# Age-dependent heterogeneity in the antigenic effects of mutations to influenza hemagglutinin

**DOI:** 10.1101/2023.12.12.571235

**Authors:** Frances C. Welsh, Rachel T. Eguia, Juhye M. Lee, Hugh K. Haddox, Jared Galloway, Nguyen Van Vinh Chau, Andrea N. Loes, John Huddleston, Timothy C. Yu, Mai Quynh Le, Nguyen T.D. Nhat, Nguyen Thi Le Thanh, Alexander L. Greninger, Helen Y. Chu, Janet A. Englund, Trevor Bedford, Frederick A. Matsen IV, Maciej F. Boni, Jesse D. Bloom

## Abstract

Human influenza virus evolves to escape neutralization by polyclonal antibodies. However, we have a limited understanding of how the antigenic effects of viral mutations vary across the human population, and how this heterogeneity affects virus evolution. Here we use deep mutational scanning to map how mutations to the hemagglutinin (HA) proteins of the A/Hong Kong/45/2019 (H3N2) and A/Perth/16/2009 (H3N2) strains affect neutralization by serum from individuals of a variety of ages. The effects of HA mutations on serum neutralization differ across age groups in ways that can be partially rationalized in terms of exposure histories. Mutations that fixed in influenza variants after 2020 cause the greatest escape from sera from younger individuals. Overall, these results demonstrate that influenza faces distinct antigenic selection regimes from different age groups, and suggest approaches to understand how this heterogeneous selection shapes viral evolution.

## INTRODUCTION

Seasonal human influenza virus undergoes continual antigenic evolution in response to immune selection from the host population. Although human individuals retain potent neutralizing antibodies to historical strains from prior exposures, they are susceptible to re-infection roughly every five years, due to rapid accumulation of mutations in the hemagglutinin (HA) surface protein that erode antibody neutralization (Kucharski et al., 2015; Ranjeva et al., 2019). Antigenically distinct strains of influenza H3N2 emerge every few years (Smith et al., 2004), resulting in frequent turnover of the virus population (Bedford et al., 2014; Fitch et al., 1997; Strelkowa and Lässig, 2012). Because of this continual antigenic evolution, individuals in the population have diverse exposure histories, which in turn shape their antibody response to subsequent infection or vaccination (Fonville et al., 2014; Francis, 1960; Krammer, 2019; Linderman et al., 2014; Skowronski et al., 2017). Neutralizing antibody titers are typically highest against strains seen early in life, and steadily decline against strains seen later in life (Fonville et al., 2014; Kucharski et al., 2015; Lessler et al., 2012; Ranjeva et al., 2019; Yang et al., 2020), which suggests that immune specificity varies, at least in part, by birth cohort.

Population-level heterogeneity has been proposed as a factor shaping the antigenic evolution of influenza (Nakajima et al., 2000; Oidtman et al., 2021; Sato et al., 2004), but we lack a detailed understanding of how the antigenic impacts of viral mutations differ across individuals in the population. Prior work has characterized this heterogeneity by analyzing human serum neutralization or hemagglutination-inhibition of recently circulating strains (Kim et al., 2023; Nakajima et al., 2000; Sato et al., 2004), or analyzing demographics of confirmed influenza cases over many seasons (Arevalo et al., 2020; Worby et al., 2015). Kim et al. (2023) found that individuals within the same age group have significantly correlated neutralization titers to different HA or NA clades. However, these studies have focused on susceptibility to naturally circulating strains, and cannot fully resolve how specific viral mutations affect neutralization by the polyclonal antibodies of different individuals.

Deep mutational scanning can be used to map the mutations in influenza HA that confer escape from neutralization by human sera, and has shown that single mutations can sometimes lead to large drops in neutralization (Lee et al., 2019). However, prior deep mutational scanning studies have not analyzed the antigenic effects of mutations across many individuals from different age groups. To address this gap, we here use deep mutational scanning to map how mutations to the HAs of two H3N2 vaccine strains (A/Hong Kong/45/2019 and A/Perth/16/2009) affect neutralization by sera collected from individuals of different ages during the timeframe that strain was in the vaccine. We find that the antigenic effects of mutations differ across individuals and especially age cohorts, with single mutations often causing large drops in neutralization. Differences among age cohorts are partially explained by inferred exposure history. We also find some evidence that the mutations that fix during natural influenza evolution in subsequent years often confer substantial neutralization escape from child and especially teenage sera. Overall, our work provides a deeper understanding of the antigenic selection exerted by different subsets of the population, and suggests ways in which this heterogeneity may shape influenza evolution.

## RESULTS

### A deep mutational scanning approach to map mutations to HA that affect viral neutralization by human serum

To systematically map how mutations to hemagglutinin affect neutralization by human sera, we developed deep mutational scanning libraries in the HA of A/HongKong/45/2019, which was the H3N2 component of the 2020-2021 influenza vaccine. The HA libraries were incorporated into replication-competent influenza virions, with the other viral genes derived from the lab-adapted WSN strain for biosafety reasons (**Figure 1A**). We improved upon prior influenza deep mutational scanning studies (Doud and Bloom, 2016; Lee et al., 2019) by incorporating barcodes into the viral genome, and by using a non-neutralized standard to allow for absolute quantification of neutralization by deep sequencing (Dadonaite et al., 2023) (**Figure 1A**, **Figure S1**-**S2**). We designed our libraries to include only mutations that are functionally tolerated in HA as assessed by prior deep mutational scanning and analysis of natural H3 HA sequences. We created two fully independent replicate HA libraries, each containing approximately 30,000 functional HA variants with an average of 3 mutations per gene (**Figure S3**).

**Figure 1.**
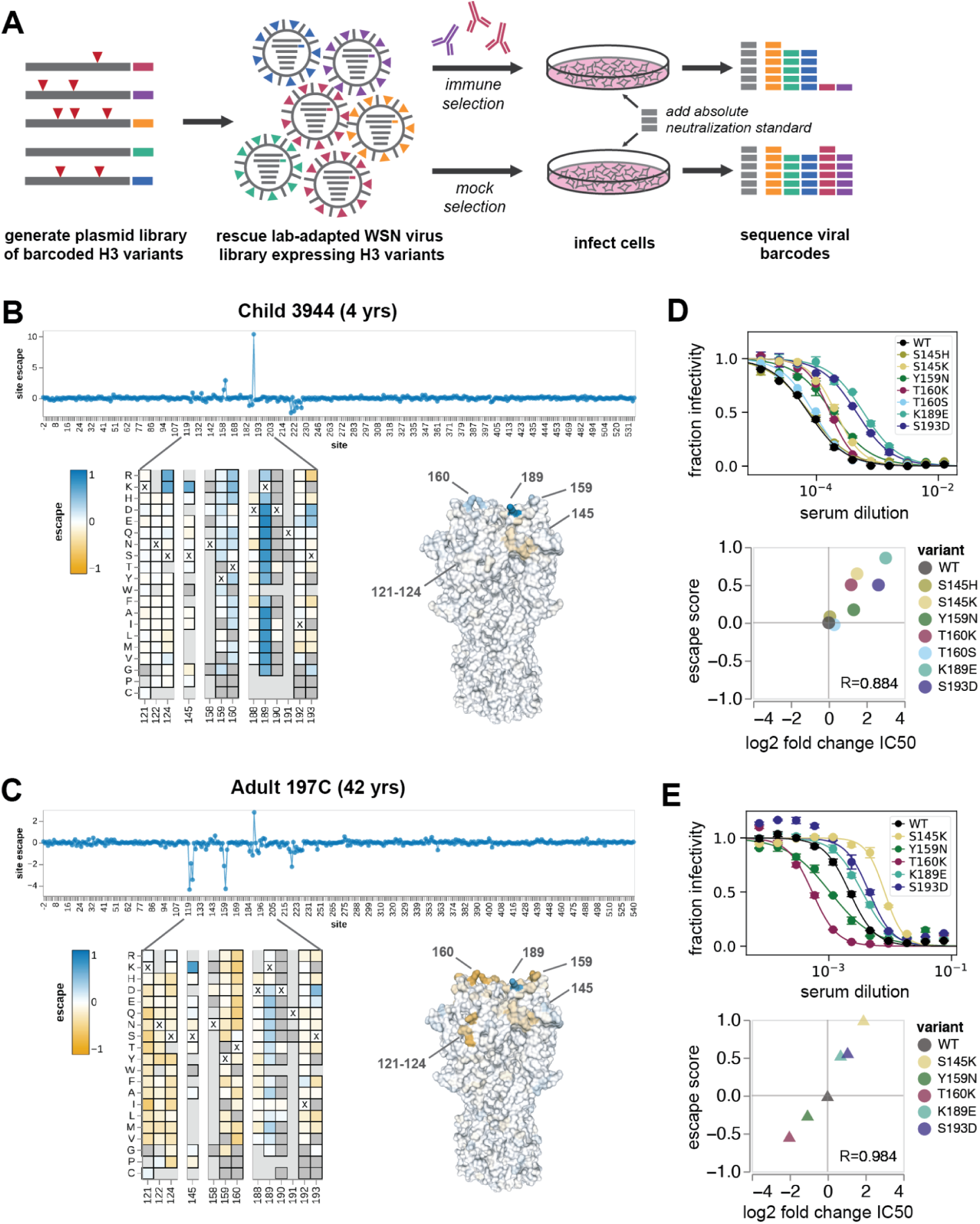
Deep mutational scanning escape maps show how mutations to HA affect viral neutralization. (A) Experimental workflow for mapping HA serum escape. The virus library is incubated with either the serum of interest, or media alone, then added to cells. At 13 hours post-infection, RNA is harvested from cells, and viral barcodes are reverse-transcribed and amplified by PCR. Escape scores are calculated for all sampled mutations by sequencing barcodes from variants that successfully infected cells in the serum-selection condition, and comparing to the mock-selection. A neutralization standard is used to convert relative sequencing counts into absolute neutralization (**Figure S2**). Example serum escape maps from a child (B) and an adult (C). The line plot shows the summed effects of all sampled mutations at each site (“site escape”, roughly proportional to fold change in neutralization), with positive values indicating neutralization escape and negative values indicating increased neutralization. Heatmaps show the effects of individual mutations for key sites. In the heatmaps, X indicates the wildtype amino acid in A/Hong Kong/45/2019 at each site, dark gray indicates mutations measured to be highly deleterious to HA-mediated infection, and light gray indicates mutations not sampled in the libraries. The structures are colored by the summed site escape scores, and show the HA structure for the A/Victoria/361/2011 H3 HA (PDB 4O5N). See https://dms-vep.github.io/flu_h3_hk19_dms/3944_escape_plot.html and https://dms-vep.github.io/flu_h3_hk19_dms/197C_escape_plot.html for full interactive escape maps with zoomable heat maps covering all sites. (D-E) Validations of escape scores by traditional neutralization assays for selected mutants against the child (D) and adult (E) sera. Correlation plots show the escape scores versus the log2 fold change in IC50 between the wildtype library strain and the mutant of interest. R indicates the Pearson correlation.

Using these deep mutational scanning libraries, we were able to map mutations across HA that increase either resistance or sensitivity of the virus to neutralization. To illustrate the type of data generated by our approach, we first show example escape maps for two sera collected in 2020 in Seattle, one from a 4-year-old child and one from a 42-year-old adult (**Figure 1B-C**). The mutation “escape scores” in these maps are roughly proportional to the log fold change in neutralization titer (Yu et al., 2022). Positive escape scores indicate mutations that confer resistance to neutralization. Negative scores indicate mutations that increase neutralization, typically by reverting sites to amino-acid identities present in older viral strains to which the individual was exposed. We validated that the escape scores measured by deep mutational scanning correlated strongly with the results of traditional neutralization assays (**Figure 1D-E**).

The escape map for the child’s serum (serum 3944) is dominated by positive escape at just a few sites, most prominently site 189 (H3 numbering), followed by sites 159 and 160 (**Figure 1B** and interactive plot linked in the figure legend). There is one region (sites 221-228) where mutations modestly increase sensitivity to neutralization. This is likely by decreasing receptor avidity rather than directly affecting antibody binding (Hensley et al., 2009), as mutations at these sites are known to alter receptor binding (Lin et al., 2012; Matrosovich et al., 2000) and have consistent neutralization-sensitizing effects across all mapped sera (**Figure S6**). At some sites (e.g. site 189), virtually any amino acid mutation causes escape, while at other sites only specific mutations cause escape (e.g. S145K) (**Figure 1B**).

In contrast, the escape map for the adult’s serum (serum 197C) reveals that some mutations increase neutralization sensitivity while others cause neutralization escape. Specifically, mutations at sites 121-124 and 159-160 increase neutralization by this serum (reflected by negative escape scores), whereas mutations at site 189 cause escape (**Figure 1C**). The sites with strongest sensitizing effects all correspond to regions where H3N2 evolution has previously introduced N-linked glycans that mask antibody epitopes. In 1997, H3N2 influenza acquired an N-linked glycan at site 122 that masked a key antigenic site. Mutations at sites 122 and 124 that eliminate this glycan greatly increase sensitivity to neutralization, suggesting that antibodies targeting this now-inaccessible site are still present at high levels in the serum. Similarly, the 158 glycan was introduced in 2014 and masked an immunodominant antigenic region (Gouma et al., 2020a; Zost et al., 2017). Increased neutralization sensitivity from mutations that remove glycosylation motifs introduced many years ago is consistent with the well-described concept of antigenic seniority, where individuals retain neutralizing antibodies against strains encountered early in life (Lessler et al., 2012).

### Antigenic effects of HA mutations differ among individuals and age cohorts

To better understand how antibody escape mutations differ among human individuals, we generated escape maps for sera collected in 2020 from 8-10 individuals each in three different age cohorts: children (2-5 years of age), teenagers (15-20 years), and adults (40-45 years) (**Figure 2**). Most individuals were vaccinated within the past year (**Tables S1-S2**). We also mapped serum from an unvaccinated 2-month old infant and a 68-year-old adult. Sera were chosen from a larger set, based on having good neutralization activity against the parental A/Hong Kong/45/2019 library strain HA (**Figure S4**). To confirm that our escape maps are replicable, we mapped two sets of sera sampled from the same individual on different days, and found that the results were almost identical (**Figure S5**).

**Figure 2.**
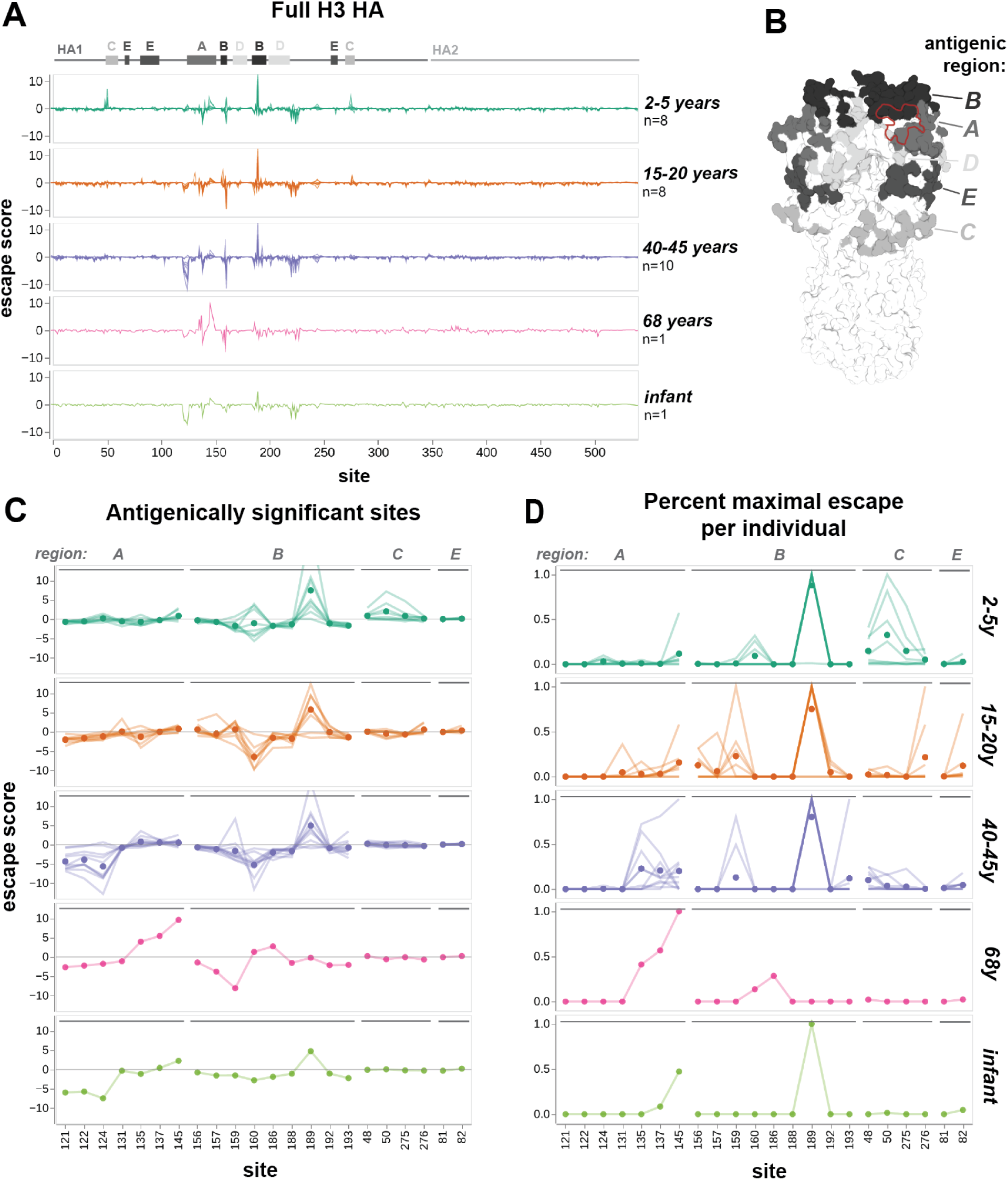
Escape maps from different cohorts show age-dependent trends in serum targeting of A/Hong Kong/45/2019HA. (A) Serum escape maps for different human individuals in each age cohort. Each line represents an individual, and individuals within the same cohort are overlaid in the same plot. Escape scores are the summed effects of all sampled mutations at each site. Approximate locations of HA1, HA2, and antigenic regions A through E (as defined by Muñoz and Deem, 2005) are labeled above the plots. (B) H3 HA trimer colored by antigenic region, with RBS outlined in red. Structure is from A/Victoria/361/2011 H3 HA (PDB 4O5N). (C) Serum escape maps at antigenically significant sites for human individuals in each age cohort. Each line is an individual escape map, and points represent the mean escape score for that age cohort at that site. The y-axis is clipped to 12.5 in both (A) and (C), although one child has an escape score of 23.9 at site 189. (D) Escape maps for key sites with escape floored at zero and normalized to the maximal positive escape score for that individual. Lines represent individuals and points represent the mean normalized escape score for that age cohort.

For all sera, the strongest escape mutations are within a small subset of sites in HA1 that correspond to classically defined antigenic regions (Wiley et al., 1981) (**Figure 2A-B**) . The mutations that cause the greatest escape are in antigenic regions A and B, which are near the sialic-acid receptor binding pocket and have alternated in immunodominance throughout H3N2 evolution (Popova et al., 2012). Mutations in antigenic regions C and E, which are lower on the HA head, cause weaker escape from the sera of some individuals. Two outlier sera, child 4584 and teenager 3856, exclusively target regions C and E and have no escape mutations in the immunodominant regions A and B (**Figure S6**). Notably, out of all mapped sera, these two sera had the lowest neutralization activity against the parental library strain.

There are clear differences in escape maps among age cohorts, though they share some sites of strong escape. This can best be seen by analyzing both the overall magnitude of escape across sera (**Figure 2C**), and the sites that confer the strongest escape for each serum, normalizing by their maximum escape score (**Figure 2D**). For most cohorts, the highest overall escape is at site 189 in the immunodominant antigenic region B. Most adult sera are also escaped by mutations in region A, which was immunodominant in the early 1990s (Nobusawa et al., 2012). Teenage sera are more often escaped by mutations at sites 156-159 in region B, which has likely been immunodominant for their entire lifetime (Chambers et al., 2015; Popova et al., 2012) (**Figure 2D**). Some child sera target a different site in region B, site 160, and additional sites in region C. These region C sites are distant from the receptor-binding pocket (**Figure 2B**), and proximal to the vestigial esterase subdomain, a target of some broadly neutralizing antibodies (Wu and Wilson, 2017). The escape map for the serum from an elderly individual is distinct from other cohorts, with the strongest escape at site 145, and no signal at 189. This individual was born 16 years before the introduction of H3N2, and differences in childhood HA exposure could contribute to their unique escape map.

Sites where mutations strongly increase neutralization sensitivity (indicated by negative values in the escape maps) are more common in older age cohorts (**Figure 2C**). There are two regions of strongly sensitizing mutations in adults: sites 121-124 and site 160. For teenagers, only site 160 exhibits strongly sensitizing mutations. Some children have weakly sensitizing mutations at site 160. Sites 158-160 and 122-124 are glycosylation motifs that appeared during the lifetimes of certain cohorts, and individuals with sensitizing mutations at these sites were exposed to prior strains without this glycan. The 122-124 glycan appeared in 1997. Adults exposed to H3N2 prior to 1997 could have generated neutralizing antibodies targeting this region, but the epitope has always been masked in strains seen by younger cohorts. The 158-160 glycan appeared in 2014, during the lifetime of both teenagers and adults, so these cohorts have similar sensitizing mutations at site 160. The weak negative signal in some children may be attributed to vaccination against A/Kansas/14/2017 H3N2 in the previous year, which does not express the 158-160 glycan (Gouma et al., 2020b) (**Table S1**). There are additional sites of weak negative signal, such as 159 and 193 in adults, and 135 in teenagers, that correspond to sites mutated during their lifetimes (**Figure S8**).

By visual inspection, it appears that teenagers and adults often have more heterogeneous escape maps within their age cohorts than children (**Figure 2D**). The main escape site for all children is site 189, with the exception of one outlier that exclusively targets antigenic region C. Secondary escape sites are limited and highly consistent among individuals. While 189 is also a major target for most teenagers and adults, the magnitude is comparable to escape at other sites for many individuals, and these other sites are variable between sera. This suggests that a naive population with limited, likely common prior exposure to influenza develops a narrow and fairly stereotyped antibody response. As exposure history increases, antibody responses diversify, even within an age cohort.

The escape map for the 2-month-old unvaccinated infant is similar to the adults, with neutralization-sensitizing mutations at sites 121-124 (**Figure 2C**). Influenza antibodies are transferred from mother to fetus during pregnancy, and confer passive immunity in newborns for several months (Nunes et al., 2015). We hypothesize that this infant gained antibodies targeting the glycan-masked 121-124 region from their mother, alongside antibodies with neutralization activity at sites 145 and 189.

In short, these escape maps demonstrate that sera from different age cohorts are escaped by mutations at different sites in HA. Children with limited prior exposure tend to have serum that is escaped by a fairly homogenous and focused set of mutations, whereas escape mutations become more heterogeneous with age. The prevalence of neutralization sensitizing mutations in the escape maps for adults indicated that older individuals often retain antibodies that target strains seen earlier in their lives. At a population level, site 189 is the most potent target of neutralizing antibodies, but individual teenagers and adults also target diverse additional sites in several known antigenic regions. This heterogeneity is especially notable given that most individuals were vaccinated against the same strains in the prior season (**Tables S1-S2**).

### Validation of the deep mutational scanning in traditional neutralization assays

We validated key mutations using traditional neutralization assays for eight representative sera from different age cohorts. The changes in IC50 measured by traditional neutralization assays strongly correlate with the escape scores measured by deep mutational scanning (**Figure 3**, **Figure S7**).

**Figure 3.**
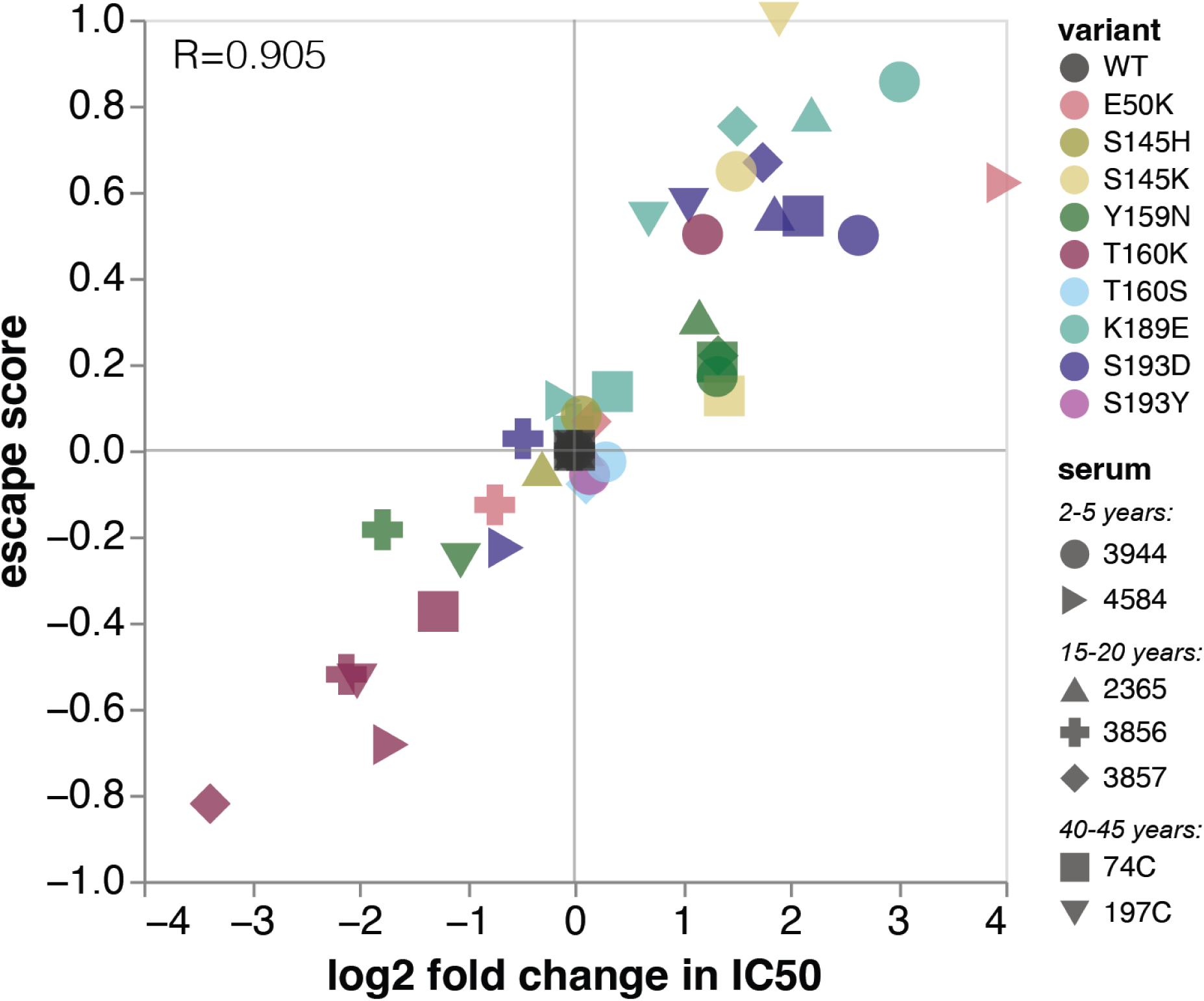
Validation of deep mutational scanning results using neutralization assays. The escape score measured by deep mutational scanning for each mutation is plotted against the log2 fold change in IC50 between the wildtype library strain and the mutant of interest. R indicates the Pearson correlation. See Figure S7 for full neutralization curves and serum-level correlation plots.

The validation neutralization assays also confirm the serum-to-serum heterogeneity observed in the deep mutational scanning. For example, Y159N confers escape from four sera, but makes the virus more sensitive to neutralization by sera 3856 and 197C (**Figure 3**). Despite K189E and S193D being strong escape mutations for almost all other sera, they had neutral or slightly sensitizing effects for sera 3856 and 4584. At some sites, only specific mutations confer escape. We confirmed that S145K and S193D were escape mutants, but S145H and S193Y were indeed neutral. These validations emphasize that antigenic effects of mutations are often highly variable across a population.

Mutations at site 160 are sensitizing for most sera, but confer escape in three children. This is especially notable because 160T is part of a glycosylation motif. It’s unclear whether these children have antibodies targeting the glycan itself, or a glycan-adjacent region, but we validated that T160K is indeed an escape mutant for child 3944. We also confirmed that T160S, which maintains the glycan, is neutral.

### Age-dependent heterogeneity in serum escape is also seen in an unvaccinated cohort measured against a different HA strain

The aforementioned trends were all based on escape maps for the HA of the A/Hong Kong/45/2019 (H3N2) strain generated using sera collected in 2020 in Seattle, Washington, USA. To assess whether trends in age-dependent heterogeneity in escape mutations are a general phenomenon, we also analyzed sera from nine children (2-4 years of age) and three adults (30-35 years of age) collected between 2010 and 2011 from Ho Chi Minh City, Vietnam against the HA of the A/Perth/16/2009 (H3N2) strain. In contrast with the high vaccination rates in the Seattle cohort (**Tables S1-S2**), vaccination rates in Vietnam are negligible (Nhat et al., 2017). We also generated escape maps for four ferrets infected with the A/Perth/16/2009 library strain. The A/Perth/16/2009 HA deep mutational scanning used an older approach with previously described libraries (Lee et al., 2019, 2018) that are not barcoded. This older deep mutational scanning approach measures relative escape scores for each serum, but the magnitude of these escape scores is not directly comparable between sera. Furthermore, the older deep mutational scanning approach can only reliably measure escape mutations, not sensitizing mutations.

The ferret and child sera have similar escape mutations within their respective cohorts, while the adult sera target heterogeneous sites (**Figure 4A-D**). All four singly-infected ferrets have almost identical escape maps and target sites 189 and 193. The children have slightly more variable escape maps, but are still quite similar to each other and all target some combination of sites 189, 193, and 159-160. In contrast, each adult serum most strongly targets a different site: 159, 160, 189, or 192. These sites are proximal to one another (**Figure 4B**), but only confer escape from certain sera, suggesting that neutralizing activity against HA from polyclonal sera is highly focused on specific residues. We validated the antigenic effects of key mutants in neutralization assays, and found that they correlated very well with the predicted escape scores (**Figure 4E**, **Figure S10**). Note that the slope of the correlation line between escape score and measured fold-change in IC50 varies among sera in **Figure 4E**, because the magnitude of escape scores for this older deep mutational scanning approach are not directly comparable among sera.

**Figure 4.**
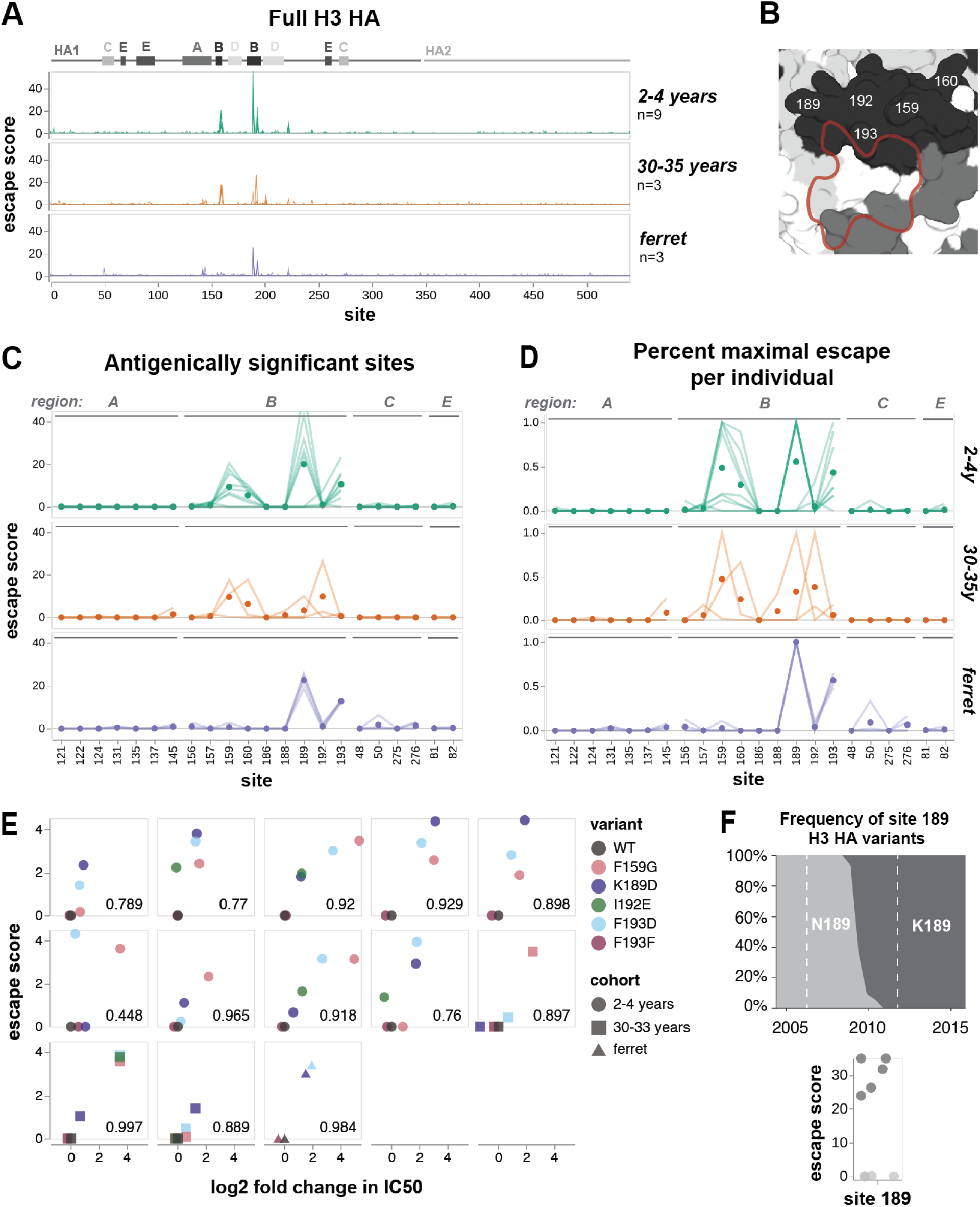
Escape maps from A/Perth/16/2009 H3 HA, measured against sera from an unvaccinated cohort and singly-infected ferrets. (A) Serum escape maps for different individuals in each cohort. Each line represents an individual, and individuals within the same cohort are overlaid in the same plot. Escape scores are the summed effects of all sampled mutations at each site. Approximate locations of HA1, HA2, and antigenic regions A through E are labeled above the plots. (B) Antigenic region B of H3 HA is shown in black, with RBS outlined in red. Structure is from A/Victoria/361/2011 H3 HA (PDB 4O5N). (C and D) Serum escape maps at antigenically significant sites for individuals in each cohort. Each line is an individual escape map, and points represent the mean escape score for the cohort at that site. (C) shows the escape scores, while (D) shows the max normalized escape scores for each individual. (E) Correlation of escape scores measured in deep mutational scanning versus the fold-change in IC50 measured in traditional neutralization assays for each serum. F193F is a synonymous mutant used as a control. The Pearson correlation is indicated for each serum. See **Figure S10** for full neutralization curves. (F) Global frequency of site 189 H3 HA variants, compared to escape scores at site 189 from 2-4-year-old children. Dashed lines on frequency plot indicate maximum exposure period for this cohort, from the earliest birth date to latest serum collection date. Frequency plot adapted from the Nextstrain real-time pathogen evolution website (Hadfield et al., 2018; Neher and Bedford, 2015).

Site 189 has high escape scores for five of the child sera, and is neutral for the other four (**Figure 4F**). The difference between these two groups of children is likely due to the arrival of the N189K mutation in H3N2 HA in 2009, which was halfway between their birth and serum collection. Children with positive escape scores at site 189 have neutralizing antibodies that can bind to K189 in the parental library strain, which suggests that they were exposed to a H3N2 virus with K at site 189. In contrast, we speculate that the children who do not have positive escape scores were likely exposed to H3N2 viruses with N189, and so do not have antibodies targeting K189 in the library strain. Note that the children with the strongest escape at sites 159-160 are all unaffected by mutations at site 189 (**Figure S9**).

Ferret sera have escape maps similar to children, but not adults (**Figure 4A,C,D**), which suggests that exposure history may be more important than species in shaping specificity. Previous work has shown that singly-infected children and ferrets have similar immune specificity, based on HAI titers against different H3N2 strains (Fonville et al., 2016). Our escape maps confirm that children and ferrets generate neutralizing antibodies against similar sites.

### Shifts in immune specificity between 2010-2011 and 2020 cohorts reflect H3N2 evolution during this time period

All the 2010-2011 sera mapped against A/Perth/16/2009 are escaped mainly by mutations in antigenic region B, whereas the 2020 sera mapped against A/Hong Kong/45/2019 are escaped by mutations in antigenic regions A, B, C, and E to varying degrees (**Figure 5A-B**). This shift in the location of key escape mutations is likely due to the introduction of the 158 glycan in 2014, which masked nearby sites in antigenic region B (**Figure 5B**) (Gouma et al., 2020a). The addition of this 158 glycan may explain why the dominant escape site in A/Hong Kong/45/2019 is 189 for all age cohorts: site 189 is still in the immunodominant antigenic region B, but is further from the 158 glycan, and has not been substituted in H3N2 evolution since 2009 (**Figure 4F**, **Figure 5B**). A/Hong Kong/45/2019 has also lost glycans at sites 133 and 144 relative to A/Perth/16/2009, which may contribute to increased escape in antigenic region A in A/Hong Kong/45/2019 by unmasking this region (**Figure 5B**).

**Figure 5.**
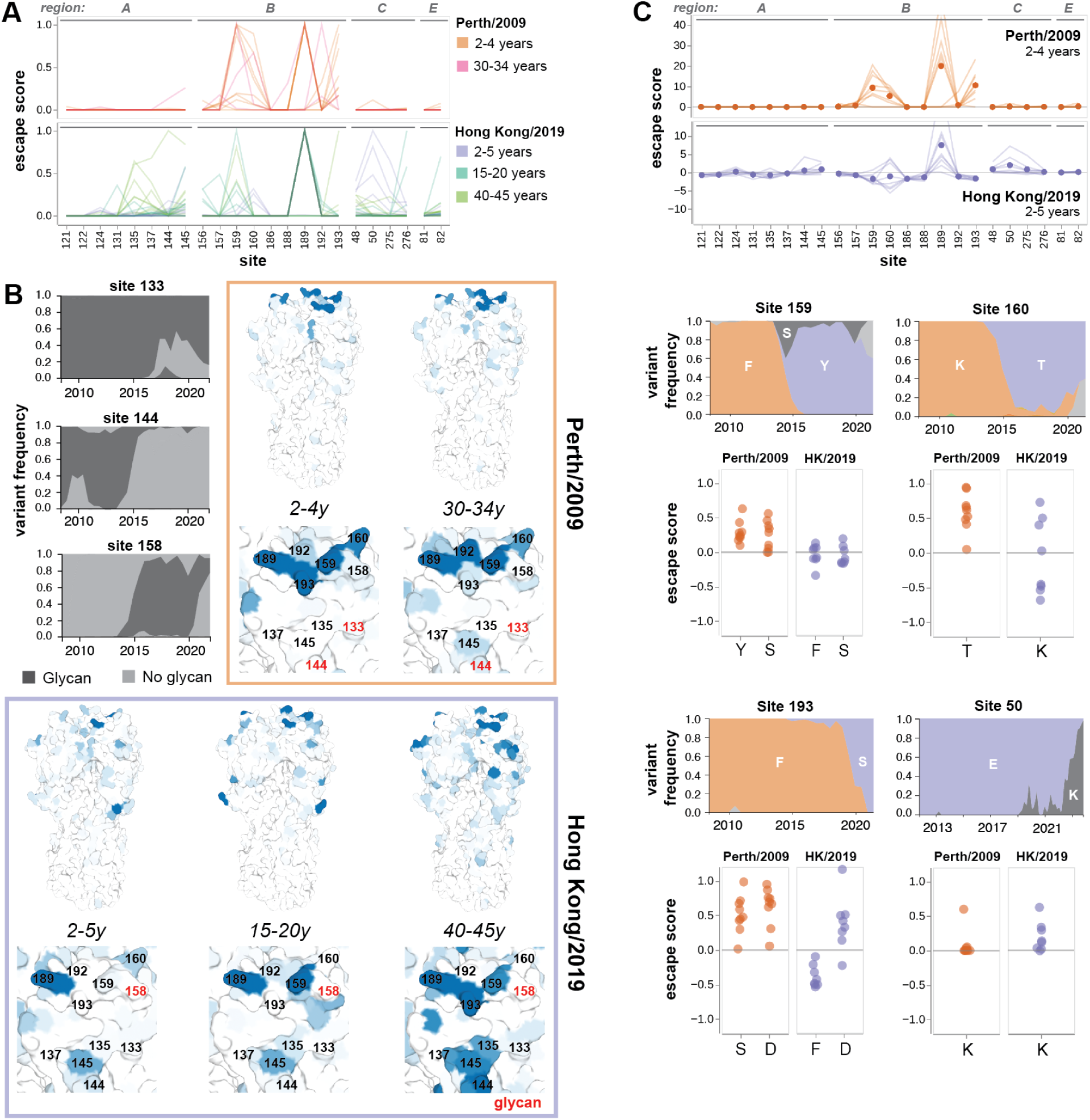
Comparison of serum escape maps between 2010-2011 and 2020 cohorts. (A) Normalized serum escape maps at antigenically significant sites for different individuals in each cohort, colored by age group. Each line is an individual escape map, normalized by the maximum escape for that individual. (B) Maximal normalized escape score for each age cohort visualized on the H3 HA structure. Zoomed panels show key sites in antigenic regions A and B. Glycosylated sites (site 133 and 144 for A/Perth/16/2009, and site 158 for A/Hong Kong/45/2019) are indicated with red text, and the global frequency of variants with these glycans over time are plotted to the left. Structures are from A/Victoria/361/2011 H3 HA (PDB 4O5N). (C) Changes in escape maps from young children between 2010 and 2020, compared to the emergence of new variants at those sites. Global frequency plots of H3 HA variants at sites 159, 160, and 193 are colored by the wildtype amino acid in A/Perth/16/2009 (F159, K160, F193) or A/Hong Kong/45/2019 (Y159, T160, S193). At site 50, both library strains have the same genotype (E50). Escape scores for mutations to amino acids circulating from 2010-2020 are plotted below. Points represent the mutation escape score for each individual in the cohort. For the 2010-2011 cohort, escape scores were normalized to one at each site, to facilitate comparison to escape scores from the 2020 cohort. Frequency plots in (B) and (C) were adapted from the Nextstrain real-time pathogen evolution website (Hadfield et al., 2018; Neher and Bedford, 2015).

Most escape maps from child sera are dominated by sites 159, 160, 189, and 193 in A/Perth/16/2009. Notably, mutations at three of these four sites rose to at least 90% frequency in the next 5-10 years (**Figure 5C**). Some children in the 2020 cohort also have sera that are escaped by mutations at site 50. The mutation E50K began to appear in circulating viruses in 2019, and reached 90% frequency in 2023 (**Figure 5C**). These evolutionary patterns suggest that the escape sites mapped here, especially from child sera, reflect antigenic pressure on influenza virus.

Site 189 in A/Hong Kong/45/2019 is generally the dominant escape site for the child sera collected in 2020 (**Figure 5C**). The differences in escape sites in the 2020 versus 2010-2011 child sera can be explained by mutations introduced between 2010 and 2020. The 158-160 glycan likely hindered the generation of antibodies targeting 159-160. However, mutations at site 160 in A/Hong Kong/45/2019 are either moderate escape or sensitizing mutations against child sera. This indicates that site 160 is still somewhat of a target of neutralizing antibodies for children in the 2020 cohort, but these antibodies overall make a smaller contribution to serum activity than for children in the 2010-2011 cohort.

Similarly, the mutation F193S was introduced in 2019, and children from the 2020 cohort were likely exposed to strains with the prior genotype F193. Reverting to F193 makes A/Hong Kong/45/2019 more sensitive to neutralization by these sera, and 193D remains a strong escape mutation for both cohorts, which indicates that these children have neutralizing antibodies against site 193 in their serum repertoire (**Figure 5C**). In short, the neutralizing antibody response of children with limited exposure history is largely directed towards sites 160, 189, and 193 in both cohorts.

### H3 HA evolution after 2020 correlates more with immune pressure imposed by children and teenagers than adults

These results show that influenza faces somewhat distinct antigenic selection regimes from different age groups. This raises the possibility that some mutations could be especially beneficial for the virus within subsets of the population (**Figure 6A**). In order to see if the actual evolution of the virus strongly favored escape mutations from specific age cohorts, we analyzed serum escape for approximately 1,200 representative H3N2 strains circulating from 2012 to 2023. The overall escape score for a given strain was calculated as the sum of the escape scores for all of its HA amino-acid mutations relative to A/Hong Kong/45/2019. For each serum, strains with negative scores are more potently neutralized than A/Hong Kong/45/2019, while strains with positive scores can better escape neutralization.

**Figure 6.**
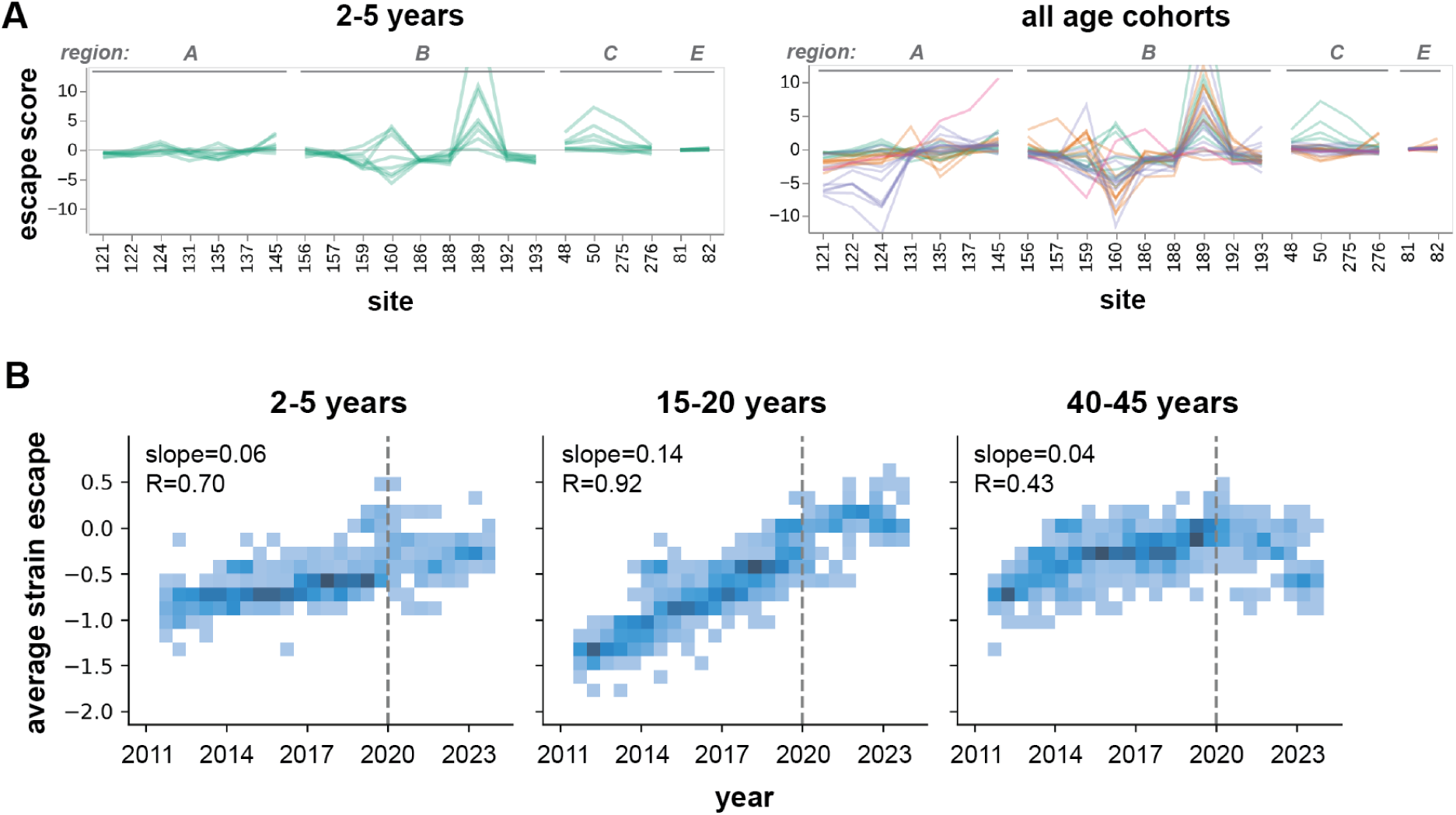
Relationship between escape mapped in deep mutational scanning and actual H3N2 HA evolution in humans. (A) Serum escape maps at key antigenic sites for individuals in either the 2-5-year-old cohort, or all age cohorts, tested against the A/Hong Kong/45/2019 HA. Each line is an individual escape map. (B) Escape scores for H3N2 influenza strains circulating from 2012 to 2023, averaged across sera within each cohort. Strain escape is calculated based on the sum of the cohort-average escape scores for all mutations in that strain, relative to the parental library strain A/Hong Kong/45/2019, which has an escape score of zero. Plots show a 2D histogram of escape scores for approximately 1,200 H3N2 HA variants sampled during this timeframe. Dashed line indicates the year of serum collection (2020). The slope of the best-fit line relating year to average strain escape (line not shown) is reported for each cohort, along with the Pearson correlation. The slope is significantly higher for the teenage cohort compared to the child (*p*=0.022) and adult (*p*=0.004) cohorts.

Analysis of the average strain escape scores from each cohort shows that the evolutionary trajectory of H3N2 during this timeframe strongly correlates with escape from teenage sera (slope=0.14, Pearson *r*=0.92) (**Figure 6B**). Escape from child sera also correlates to a lesser degree (slope=0.06, r=0.70), although this timeframe includes years prior to their birth. In contrast, mutations gained after 2020 did not substantially escape neutralization by serum from the adult cohort. We confirmed that the increase over time in average strain escape scores from teenage sera are significantly higher than adult sera (*p*=0.004) or child sera (*p*=0.022). Although the number of sera in our analysis is modest, these results show how different age cohorts are differentially affected by viral evolution, and suggest that teenagers and children could play an especially prominent role in driving viral evolution.

## DISCUSSION

We have mapped the mutations that escape neutralization by the polyclonal serum antibodies of 40 human individuals from different age cohorts. Sites of neutralization escape for any given serum are often highly focused, with a few specific mutations often leading to 5-10 fold drops in neutralization by polyclonal serum antibodies—a finding consistent with prior work (Davis et al., 2018; Huang et al., 2015; Lee et al., 2019; Linderman et al., 2014). We also identify mutations that increase virus sensitivity to neutralization, typically by restoring epitopes present in older strains, such as removal of recently acquired N-linked glycosylation sites. The existence of these sensitizing mutations likely constrains the virus’ evolution by disfavoring reversions to epitopes present in older strains.

The effects of mutations on neutralization escape differ between age cohorts in a manner consistent with likely exposure history. Many teenage sera are most escaped by mutations in antigenic region B, whereas adult sera are more strongly affected by mutations in antigenic region A, which reflects the regions that were likely immunodominant during their respective childhoods (Chambers et al., 2015; Nobusawa et al., 2012; Popova et al., 2012). Heterogeneity between cohorts can be highly specific: although some child and teenage sera are escaped by mutations in antigenic region C, these mutations are at site 50 for children and site 276 for teenagers. The relative number and magnitude of neutralization-sensitizing mutations also increases with age, likely due to older individuals having more antibodies from prior exposures to older strains.

The child sera analyzed in this study are escaped by mutations at a fairly narrow and homogenous set of sites. This homogeneity suggests a relatively stereotyped initial neutralizing antibody response focused on sites near the receptor binding pocket of HA, which is consistent with prior characterization of the antibody response in children (Islam et al., 2017; Meade et al., 2020; Nakajima et al., 2000). Four out of five sites targeted by children have mutated in subsequent years, suggesting that deep mutational scanning can be used to identify HA sites under strong immune pressure in contemporary human influenza strains.

Exposure history may be more important than species in shaping neutralizing specificity, as ferrets infected with A/Perth/16/2009 have similar escape mutations to children from the 2010-2011 cohort. The similarities between ferrets and naive children, but not adults with complex exposure histories, further illustrates the limitations of using primary ferret antisera as a proxy for adult human sera (Fonville et al., 2014; Lee et al., 2019; Linderman et al., 2014; Li et al., 2013). Nevertheless, ferret sera may still be useful for identifying the selective pressures imposed by young children (Fonville et al., 2016).

The introduction of the 158-160 glycan on the HA head in 2014 resulted in major antigenic change, as it hindered antibody binding to the immunodominant antigenic region B, leaving many individuals highly vulnerable to infection (Gouma et al., 2020a; Zost et al., 2017). It was therefore surprising that this glycan was subsequently lost in 2021 with the mutation T160I (Bolton et al., 2022). Our analysis of sera from 2020 identifies three children whose serum neutralization is escaped by mutations at site 160, suggesting that the T160I mutation may have been gained in response to antigenic selection from young children.

Our data illustrate how mutations to HA can have heterogeneous effects on virus neutralization by sera from different individuals in the human population. The heterogeneity in escape mutations is substantially clustered by age cohort, raising the question of whether certain age groups (e.g., children, teenagers, or adults) might play an especially important role in driving viral antigenic evolution (Kim et al., 2023; Nakajima et al., 2000). Previous work has proposed that school-aged children are important drivers of influenza epidemics in communities, based on transmission modeling (Basta et al., 2009; Wallinga et al., 2006), earlier peaks of infection in this age group (Worby et al., 2015), and the impact of school closure on transmission (Cauchemez et al., 2008; Huang et al., 2014). Whether the populations driving epidemics also drive antigenic drift is currently unclear. Although our study does not examine a sufficient number of sera or viruses to draw definitive conclusions, our results suggest that children and teenagers play an important role in shaping viral evolution, since mutations that fixed in HA often cause more escape from child and teenage sera than adults. More extensive characterization of the major escape mutations in different age cohorts and viruses could further test this hypothesis, and any insights into which segments of the population drive antigenic evolution could be used to improve evolutionary forecasting for influenza vaccine-strain selection.

Heterogeneity in the antigenic effects of mutations across the population could also have an effect on the overall rate of evolution of human influenza virus. If a specific mutation confers escape from neutralizing antibodies across all individuals, it will have a greater beneficial effect than if it only confers escape from some individuals—and this heterogeneity could impact evolutionary dynamics (Cobey and Pascual, 2011; Desai and Fisher, 2007; Gupta and Galvani, 1999; Luksza and Lässig, 2014). We therefore suggest that continued characterization of population-level heterogeneity in viral escape mutations, and incorporation of these measurements into models of evolution, could shed further light on the evolutionary dynamics of influenza and other viruses.

### Limitations of this study

Our experiments analyzed sera from a limited number of individuals from each age cohort, and these cohorts were sampled from a single city for each year (Seattle for the 2020 cohort, and Ho Chi Minh City for the 2010-2011 cohort). While this sample size was sufficient to illustrate heterogeneity in serum antibody targeting between age cohorts, larger sample sizes are needed to draw firm conclusions about how the heterogeneity in the antigenic effects of mutations shapes influenza evolution.

A further caveat is that we focused on sera that had high neutralizing titers against the vaccine strains used to build our libraries, in large part to reduce the serum volumes needed to perform the deep mutational scans. Therefore, if the antigenic effects of mutations vary systematically with the overall serum neutralizing titer, then our results might only reflect the subset of the population with high titers.

Our study coincides with the beginning of the SARS-CoV-2 pandemic, which may limit the generalizability of results measured from the 2020 cohort. More specifically, differences in social distancing measures implemented for children and adults may have influenced which age groups were stronger drivers of influenza evolution in subsequent years.

## Acknowledgements

We thank Sarah Cobey for scientific advice. We gratefully acknowledge all data contributors, including the authors and their originating laboratories responsible for obtaining the specimens, and their submitting laboratories for generating the genetic sequence and metadata and sharing via the GISAID Initiative, on which part of this research is based. This work was supported in part by the NIH/NIAID award R01AI165821 to TB and JDB, contract 75N93021C00015 to JDB and TB, and award R01AI146028 to FAM. Serum collection in Vietnam was funded by a Wellcome Trust/Royal Society Sir Henry Dale Fellowship (098511/Z/12/Z) to MFB. TCY was supported by the CMB training grant T32 GM007270 and the NSF graduate research fellowship DGE-2140004. JDB, TB, and FAM are Investigators of the Howard Hughes Medical Institute. This research was supported by the Genomics & Bioinformatics Shared Resource (RRID:SCR_022606) of the Fred Hutch/University of Washington/Seattle Children’s Cancer Consortium (P30CA015704), and by Fred Hutch Scientific Computing (NIH grants S10-OD-020069 and S10-OD-028685).

## METHODS

### Data Availability Statement

The file https://github.com/dms-vep/flu_h3_hk19_dms/blob/main/results/full_hk19_escape_scores.csv contains the fully processed serum escape data for sera from the 2020 cohort measured against the A/Hong Kong/45/2019 library. This file reports the mean escape score for each mutation, averaged between the two replicate libraries. Entries are filtered to mutations that are represented in at least three independent variants, are not stop codons, and do not have highly deleterious functional effects. Functional effects represent the effect of a mutation on HA protein function, calculated based on the ratio of mutation counts in the plasmid library compared to the passaged virus library. This file only includes mutations with functional effects greater than -1.38. Sites are numbered using standard H3 numbering, where the first codon of the ectodomain is site 1, through the HA1 subdomain. Rather than switching to (HA2)1 at the beginning of the HA2 subdomain, as is standard, we then continue with sequential numbering (i.e. site (HA2)1 in standard H3 numbering corresponds to site 330 in this data).

The file https://github.com/dms-vep/flu_h3_hk19_dms/blob/main/results/perth2009/merged_escape.csv contains the fully processed serum escape data for sera from the 2010-2011 cohort measured against the A/Perth/16/2009 library. This file reports the median escape score for each mutation from three replicate library measurements. Sites are numbered using standard H3 numbering.

See https://github.com/dms-vep/flu_h3_hk19_dms for the full analysis pipeline for deep mutational scanning of A/Hong Kong/45/2019, and https://dms-vep.github.io/flu_h3_hk19_dms/ for interactive documentation of the results. See https://github.com/dms-vep/flu_h3_hk19_dms/tree/main/figures for code used to generate all main figures in this paper, with the exception of Figure 6B, which can be found at https://github.com/matsengrp/seasonal-flu-dmsa (described in the Computational Methods section ‘Estimating escape scores for circulating H3N2 strains’).

See https://github.com/jbloomlab/map_flu_serum_Vietnam_H3_Perth2009 for the full analysis of deep mutational scanning of A/Perth/16/2009.

### Biosafety

All mutant viruses in the deep mutational scanning studies derived their HAs from one of two recent human H3N2 vaccine strains (A/Perth/16/2009 or A/Hong Kong/45/2019) and the remaining seven genes from the lab-adapted A/WSN/1933 (H1N1) strain. These viruses are therefore classified as biosafety-level 2, and the work was approved by the Fred Hutchinson Cancer Center biosafety committee.

### Human sera

The human sera used for mapping escape from A/Hong Kong/45/2019 were collected in Seattle, Washington in 2020. The sera from the 2-month-old infant, the 2-5-year age cohort, and the 15-20-year age cohort are residual sera collected at Seattle Children’s Hospital with approval from the Human Subjects Institutional Review Board. Sera from the 40-45-year age cohort were collected as part of the prospective longitudinal Hospitalized or Ambulatory Adults with Respiratory Viral Infections (HAARVI) cohort study of individuals with SARS-CoV-2 infection. Written informed consent was obtained for each participant. Serum from the 68-year-old individual is deidentified residual serum from the University of Washington Virology Lab that was collected for testing for HBsAb (for which they tested negative), under a study approved by the University of Washington IRB with a consent waiver.

The human sera used for mapping escape from A/Perth/16/2009 were obtained from a serum bank established in 2010 in central and southern Vietnam for the purposes of calculating disease burden, pathogen exposure, and vaccine coverage for a range of pathogens (Berto et al., 2018; Boni et al., 2013; Lam et al., 2019; Nhat et al., 2017; Quan et al., 2018; Thwaites et al., 2023; Vinh et al., 2021). Anonymized samples were collected, as residual serum, from the biochemistry and haematology labs of the Hospital for Tropical Diseases in Ho Chi Minh City, Vietnam, during 2010 and 2011. The study was approved by the Scientific and Ethical Committee of the Hospital for Tropical Diseases and the Oxford Tropical Research Ethics Committee at the University of Oxford.

### A/Perth/16/2009 and A/Hong Kong/45/2019 library overview

The analysis in this paper uses two distinct libraries: a non-barcoded library in the background of A/Perth/16/2009, and a barcoded library in the background of A/Hong Kong/45/2019. Because the A/Perth/16/2009 library is non-barcoded, these selection experiments use a barcoded-subamplicon sequencing approach, where the HA gene is amplified as a collection of short, tiled fragments for Illumina sequencing (see (Doud and Bloom, 2016) and https://jbloomlab.github.io/dms_tools2/bcsubamp.html for more detailed descriptions of this approach). There is no linkage information between these fragments, so escape scores are calculated as the enrichment of a given mutation in the serum selection relative to the mock selection. These libraries are therefore designed to be mostly single-mutant variants.

In contrast, the incorporation of barcodes in the A/Hong Kong/45/2019 libraries allows for more straightforward sequencing, and the ability to resolve multiple mutations in a single variant. These libraries were therefore designed to have an average of 2-3 mutations per variant. Escape scores are calculated as the enrichment of a given *variant* in a selection experiment (as measured by sequencing counts for each unique barcode), rather than enrichment of each single mutation.

The A/Perth/16/2009 experiments use the exact library stocks generated in (Lee et al., 2019). Serum selection experiments and data analysis using the A/Perth/16/2009 library were carried out as described in the methods of (Lee et al., 2019). The following methods sections all pertain to analyses using the A/Hong Kong/45/2019 barcoded libraries.

### Design of barcoded H3 plasmid with duplicated WSN-HA packaging signals

Influenza is multi-segmented, and proper packaging of vRNA into the virion relies on segment-specific RNA sequences called “packaging signals.” These span both coding and noncoding regions at the 3’ and 5’ ends of the vRNA segment (Li et al., 2021). To insert a barcode without disrupting vRNA packaging, we designed a chimeric construct with a duplicated 5’ packaging signal (negative sense), based on chimeric influenza genes designed by (Gao and Palese, 2009). We place the full 150-nucleotide 5’ packaging signal (negative sense) from the lab-adapted strain A/WSN/1933 HA after the stop codon of the HA gene. The 90-nucleotide native packaging sequence in the 5’ coding region is synonymously recoded to avoid interference between duplicated sequences. We found that maintaining the native H3 HA 5’ coding region was essential for virus growth, consistent with prior work (Zhang et al., 2000). The native 3’ packaging signal (defined as the 32-nucleotide noncoding region and first 16 amino acids of HA) is also replaced by the corresponding sequence from A/WSN/1933 HA (32-nucleotide noncoding region and first 19 amino acids) (Wu et al., 2017) (**Figure S1**).

An extra stop codon was added to the end of the coding sequence to minimize translational readthrough. This is followed by a random 16-nucleotide barcode, then the Illumina read1 priming sequence, then the full-length WSN-HA packaging signal. Homology between the native and duplicated packaging signal could lead to barcode loss; we therefore included a stop codon in the duplicated packaging signal that will be in-frame if it replaces the native packaging signal. This ensures that loss of the barcode will generate a truncated, non-functional HA protein (**Figure S1**).

The barcoded WSN-flanked H3 HA gene is inserted into the pHH21 plasmid, flanked by the human polymerase I promoter and the mouse polymerase I terminator (Neumann et al., 1999). See https://github.com/dms-vep/flu_h3_hk19_dms/blob/main/library_design/plasmid_maps/3023_pHH_ WSNHAflank_HK19_H3_barcoded.gb for an annotated map of the full plasmid.

### Design of barcoded H3 deep mutational scanning library

In a library of HA variants with combinations of mutations, the rate of non-functional variants will be much higher than a single-mutant library, as a variant carrying *any* deleterious mutation will likely be non-functional. We therefore maximized the number of functional variants in the library by excluding mutations with highly deleterious functional effects. Using H3 amino acid preferences calculated from previous deep mutational scanning of A/Perth/16/2009 H3 HA (Lee et al., 2018), we retained mutations with functional effects more positive than the 0.75 quantile of stop codon effects (**Figure S3A**).

We then identified excluded mutations that were tolerated in other sequence contexts by analyzing natural H3 sequences circulating from 1968 to 2021. After basic quality control and exclusion of egg-passaged strains, we aligned 7,597 H3 variants, and identified 44 excluded mutations that were present at a frequency of at least 0.5% in natural sequences. These were added back into the library, for a total of 2,505 targeted mutations (**Figure S3C**).

To ensure that antigenically significant mutations were highly represented in multiple backgrounds, we biased mutagenesis towards 22 selected epitope sites (**Figure S3B**). These were defined as sites that had been mutated in at least one major clade from 2015-2021. These sites would be randomly mutated, regardless of functional effects, at a higher rate than the general targeted mutations.

Sites within a primer length of one another (approximately 30bp) will not be mutated in the same variant, because a primer introducing a mutation at position 1 carries the wildtype sequence at position 2. We therefore generated paired epitope primers for epitope sites within a primer length of one another, to ensure that major epitope sites are represented in combination.

Primers were designed to introduce these targeted mutations into the HA of A/Hong Kong/45/2019, and generated as both a forward and reverse primer pool. We generated a separate set of NNK primers to introduce random mutations at the key epitope positions, and mixed them with the corresponding forward or reverse primer pool at a 10:1:1 molar ratio of epitope : paired epitope : general primers.

See https://github.com/dms-vep/flu_h3_hk19_dms/tree/main/library_design for the full analysis described here, including identification of deleterious mutations, analysis of natural H3 sequences, identification of key epitope positions, and generation of the complete set of forward and reverse mutagenic primers.

### Generation of plasmid library of barcoded H3 HA variants

See https://github.com/jbloomlab/CodonTilingPrimers for a general description of the PCR mutagenesis strategy we use here. Note that we are using primers that generate targeted amino-acid mutations, as described above.

To mutagenize the A/Hong Kong/45/2019 HA sequence, we first amplified the HA coding sequence from the plasmid containing the barcoded WSN-flanked H3 HA gene under control of the pHH21 promoter. See https://github.com/dms-vep/flu_h3_hk19_dms/blob/main/library_design/plasmid_maps/3023_pHH_ <u>WSNHAflank_HK19_H3_barcoded.gb</u> for the sequence of this plasmid. The PCR was performed with the following conditions: PCR mix: 20µL H2O, 1.5µL 10µM forward linearizing primer (primer_088), 1.5µL 10µM reverse linearizing primer (primer_089), 2µL 5ng/µL template plasmid, and 25µL 2x KOD Hot Start Master Mix (Sigma-Aldrich, Cat. No. 71842). Cycling conditions: (1) 95C/2min (2) 95C/20sec (3) 70C/1sec (4) 54C/30sec, cooling at 0.5C/sec (5) 70C/40sec (6) Return to Step 2 x19.

The amplified, linearized chimeric H3 HA sequence was gel purified using the NucleoSpin Gel and PCR Clean-up kit (Takara, Cat. No. 740609.5) and then purified using Ampure XP beads (Beckman Coulter, Cat. No. A63881) at 1:1 sample to bead ratio.

The linearized chimeric H3 HA sequence was then mutagenized using a modification of a previously described PCR mutagenesis technique (Bloom, 2014). Forward and reverse primers for mutagenic PCR were pooled separately at a molar ratio of 10:1:1 between epitope : paired epitope : general mutagenic primers. Mutated fragments of the chimeric H3 HA sequence were then PCR-amplified in two separate reactions, using either the forward primer pool and reverse linearizing primer, or the reverse primer pool and forward linearizing primer. We performed two rounds of mutagenesis, with 7 mutagenic PCR cycles the first round and 11 the second round. The mutagenesis PCRs were performed with the following conditions: PCR mix: 20µL H2O, 1.5µL 4.5µM forward or reverse primer pool, 1.5µL 4.5µM reverse (primer_089) or forward (primer_088) linearizing primer, 4µL 3ng/µL linearized template, and 15µL KOD. Cycling conditions: (1) 95C/2min (2) 95C/20sec (3) 70C/1sec (4) 50C/30sec, cooling at 0.5C/sec (5) 70C/55sec (6) Return to Step 2 for the number of cycles described above (7) 70C/2min. Note that mutagenesis was performed in duplicate, to generate two independently mutagenized replicate libraries.

Between each round of mutagenic PCR, the mutagenized partial H3 HA fragments from the forward and reverse reactions were joined in a separate PCR reaction to generate full-length H3 HA sequences. The joining PCR was performed with the following conditions: PCR mix: 4 µL each of 1:4 diluted forward and reverse mutagenesis reactions, 1.5µL 4.5µM forward primer (primer_088), 1.5µL 4.5µM reverse primer (primer_089), 3µL H2O, and 15µL KOD. Cycling conditions: (1) 95C/2min (2) 95C/20sec (3) 70C/1sec (4) 50C/30sec, cooling at 0.5C/sec (5) 70C/40sec (6) Return to Step 2 x19 (7) 70C/2min. Joined PCR mutagenesis products were gel and Ampure bead purified after each joining reaction, as described previously.

We then appended random 16-nucleotide barcodes to the mutagenized H3 HA sequences, followed by a constant overlap sequence for cloning into the backbone plasmid, using a PCR with the following conditions: PCR mix: 11.2µL H2O, 0.9µL 10µM forward linearizing primer (primer_088), 0.9µL 10µM reverse barcoding primer (primer_090), 2µL 15ng/µL mutagenized template, and 15µL KOD. Cycling conditions: (1) 95C/2min (2) 95C/20sec (3) 70C/1sec (4) 50C/30sec, cooling at 0.5C/sec (5) 70C/40sec (6) Return to Step 2 x9 (7) 70C/2min.

The barcoded mutagenized H3 HA sequences were gel and Ampure bead purified as described previously, using a 1:1 sample to bead ratio, then cloned into a plasmid containing the flanking WSN HA packaging signals, 5’ and 3’ NCRs, and pHH21 promoter and terminator. The plasmid contains GFP in place of the H3 HA ectodomain to ensure that virions cannot be generated from untransformed plasmid. See https://github.com/dms-vep/flu_h3_hk19_dms/blob/main/library_design/plasmid_maps/2851_pHH__WSNHAflank_GFP_H3-recipient_duppac-stop.gb for detailed map of the GFP recipient plasmid. The plasmid was digested with BsmBI and NotI, then gel and Ampure bead purified. We then used a 2:1 insert to vector ratio in a 1 hour HiFi assembly reaction using NEBuilder HiFi DNA Assembly kit (NEB, Cat. No. E2621). HiFi assembly products were Ampure bead purified and eluted into 20uL of H2O for higher electroporation efficiency.

We electroporated 20µL of 10-beta electrocompetent E. coli cells (NEB, C3020K) with 1µL of the purified HiFi product. Electroporated cells were plated on 15cm LB+ampicillin plates at an estimated bottleneck of 80,000 to 100,000 colony forming units to limit library size. After approximately 18 hours of outgrowth, colonies were scraped into liquid LB+ampicillin, and grown at 37°C for 2.5 hours in liquid culture prior to plasmid purification using QIAprep Spin Miniprep Kit (Cat. No. 27104). We generated two independently mutagenized plasmid libraries, referred to as libA and libB. The final structure of the mutagenized H3 HA library construct is illustrated in **Figure S1B**.

### PacBio sequencing of H3 HA plasmid library

We used long-read PacBio sequencing to link variants to their random nucleotide barcodes, and determine the composition of the plasmid library, similar to the approach used in (Starr et al., 2020). PacBio sequencing inserts were prepared from the purified plasmid pools via NotI-HF restriction digest, rather than PCR amplification, which eliminates the possibility of PCR strand exchange scrambling barcodes. Digested inserts were then gel and Ampure bead purified, and eluted into 30uL of EB. Each library was then barcoded for PacBio sequencing using SMRTbell prep kit 3.0, bound to polymerase using Sequel II Binding Kit 3.2, and then sequenced using a PacBio Sequel IIe sequencer with a 20-hour movie collection time. Analysis is described in the section ‘PacBio sequencing data analysis’.

### Generation of H3 HA variant library in the context of lab-adapted WSN virus

Mutant virus libraries were generated by reverse genetics (Hoffmann et al., 2000). We used 6-well plates rather than 15cm plates for transfection, to limit the extent to which jackpotting of certain variants in each well could bottleneck library diversity. We transfected 40 wells of 6-well plates for each library. Each well was plated with 4x10^5^ HEK-293T cells in D10 media (DMEM, supplemented with 10% heat-inactivated FBS, 2 mM L-glutamine, 100 U/mL penicillin, and 100 μg/mL streptomycin). 21 hours after plating, cells were transfected with 220ng each of H3 HA plasmid library (or wildtype control), the pHW18* series of plasmids (Hoffmann et al., 2000) for all non-HA viral genes, and pHAGE2-EF1aInt-TMPRSS2-IRES-mCherry (Lee et al., 2018). The pHW18* plasmids encode genes from the lab-adapted A/WSN/1933 strain, and so all genes in our viruses derive from the lab-adapted WSN strain except for the HAs. We used Bioland BioT (Cat. No. B01-01) for the transfections, and followed the manufacturer’s recommendations for the protocol and DNA / transfection reagent ratios.

At 20 hours post-transfection, we removed the D10 media and added 3x10^5^ MDCK-SIAT1-TMPRSS2 cells per well in low-serum IGM (Influenza Growth Media, consisting of Opti-MEM supplemented with 0.01% heat-inactivated FBS, 0.3% BSA, 100 U/mL penicillin, 100 μg/mL streptomycin, and 100 μg/mL calcium chloride). Note that overlay of MDCK-SIAT1-TMPRSS2 cells post-transfection yielded higher virus titers than transfection of co-cultured cells in pilot rescue experiments. At 49 hours post-transfection, the transfection supernatants were harvested, clarified by centrifugation at 2,000x*g* for 5min, aliquoted, frozen at -80°C, and titered in MDCK-SIAT1-TMPRSS2 cells. The titers were 3,162 TCID50/µL for both libA and libB.

To generate a genotype-phenotype link, where the barcoded HA gene carried by each virion matches the HA variant expressed on its surface, we passaged the libraries at a low MOI such that each cell should be infected by one virion at most. For each library, we infected cells in two 5-layer 875 cm^2^ flasks (Corning, Cat. No. 353144) at an MOI of 0.01. For each flask, we gently mixed 7.875x10^7^ MDCK-SIAT1-TMPRSS2 cells in 150mL of IGM (targeting a density of 9x10^4^ cells/cm^2^ when plated) with 7.875x10^5^ TCID50 of library transfection supernatant in a sterile 250mL bottle. The cell-virus mix was then added directly to the 5-layer flask. Virus supernatant was collected at 45 hours post-infection, clarified by centrifugation at 2,000x*g* for 5min, split into 1mL aliquots, frozen at -80°C, and titered in MDCK-SIAT1-TMPRSS2 cells. The titers were 3.09x10^4^ and 1.43x10^4^ TCID50/µL for libA and libB, respectively. Assuming a maximum library size of 100,000 variants, libraries were passaged at over 10x coverage of variant diversity.

For future deep mutational scanning studies, we would recommend rescuing the library at 40 hours post-transfection, and reducing passaging to approximately 36 hours. These conditions yielded similar titers, and minimizing library growth times will reduce bottlenecking from variants carrying tissue-culture-adaptive mutations.

### Design of external neutralization standards

In order to generate quantitative escape measurements, we designed barcoded standards that would not be neutralized by human sera (**Figure S2A**). These would be spiked into the library at a low level (approximately 1%), and used to normalize the barcode frequency between mock selections and serum selections (**Figure 1A**). This normalization is expected to correct for stochastic variation in virus replication across different conditions and experiments. As more of the library is neutralized at increasingly potent serum concentrations, the proportion of library barcodes to standard barcodes will decrease in a consistent manner (**Figure S2B**).

We included standards at two stages: infection with the virus library, and harvest of RNA from infected cells. The virus standard is identical to the WSN-flanked H3 HA library construct, but uses the ectodomain and 5’ coding region from A/Turkey/Mass/1975 H6 HA, a low-pathogenicity avian HA strain (GenBank Accession AB296072.1) (Sandbulte et al., 2009) (**Figure S2A**). Note that although the HA of this viral standard is from a low pathogenicity avian-strain, the virus itself is still only biosafety-level 2, because the seven non-HA gene segments and the H6 HA packaging signals are derived from the lab-adapted WSN strain. This standard will have similar infection dynamics as the library H3 HA variants. We tested neutralization activity of several sera against the WSN-H6 HA standard, and found that any serum used at a dilution of approximately 1:100 or higher begins to neutralize WSN-H6 HA.

Therefore, to allow for analysis of low-potency sera that must be at high concentration for selections, we also included an RNA spike-in standard. This is an RNA fragment that is identical to the WSN-flanked H3 HA sequence from the 5’ coding region to the 5’ UTR, but replaces the H3 ectodomain with GFP (**Figure S2A**). This ensures that the RNA spike-in standard is a similar length to the HA RNA. The fragment is generated with a pool of approximately 100 random barcodes. We add a standardized amount of RNA spike-in to the lysis buffer used for harvest of cellular RNA, so that it’s equally distributed between wells in each experiment. We tested a range of concentrations, and found that the RNA spike-in behaves similarly to the WSN-H6 HA standard when 0.05ng of RNA spike-in is added per well (**Figure S2C**).

See https://github.com/dms-vep/flu_h3_hk19_dms/tree/main/library_design/plasmid_maps for annotated plasmid maps of both standards.

### Generation of WSN-H6 HA neutralization standard

The WSN-H6 HA standard was generated from the same GFP recipient vector used to clone the WSN-flanked H3 HA library construct (see https://github.com/dms-vep/flu_h3_hk19_dms/blob/main/library_design/plasmid_maps/2851_pHH_WSNHAflank_GFP_H3-recipient_duppac-stop.gb for plasmid map). The plasmid was digested with BsmBI and XbaI, then gel and Ampure bead purified. To avoid sequencing errors, we designed four different barcodes for the WSN-H6 HA standard, such that the nucleotide at a given site was different in all four barcodes. The four barcoded H6 HA ectodomain sequences were synthesized from Twist with a 20bp overlap with the digested recipient vector, and cloned into the recipient vector using a 2:1 insert to vector ratio in a 1 hour HiFi assembly reaction. NEB 5α High Efficiency Cells (Cat. No. C2987) were transformed with 2µL of the HiFi assembly, and plated on LB+ampicillin for 18 hours of outgrowth. Single colonies were grown in liquid LB+ampicillin at 37°C for 18 hours, plasmid was purified using QIAprep Spin Miniprep Kit, and sequence was confirmed by Sanger sequencing. The four barcoded plasmids were then combined at equal concentrations. See plasmids 3238-3241 in https://github.com/dms-vep/flu_h3_hk19_dms/tree/main/library_design/plasmid_maps for detailed map of the WSN-H6 HA plasmid with four unique barcodes.

To generate the virus, we transfected a coculture of 4x10^5^ HEK-293T and 5x10^4^ MDCK-SIAT1 cells with 220ng each of the pooled WSN-H6 HA plasmid, the pHW18* series of plasmids (Hoffmann et al., 2000) for all non-HA viral genes, and pHAGE2-EF1aInt-TMPRSS2-IRES-mCherry. Transfections were performed in D10 media using BioT transfection reagent. Media was changed to IGM at 20 hours post-transfection. At 68 hours post-transfection, transfection supernatants were harvested, clarified by centrifugation at 2000x*g* for 5 min, aliquoted, frozen at -80°C, and titered in MDCK-SIAT1-TMPRSS2 cells. Titers were 3.16x10^5^ TCID50/µL.

### Generation of RNA spike-in neutralization standard

The RNA spike-in standard was generated from the GFP recipient vector, with the final 46 amino acids from the library strain appended at the 5’ end of the GFP sequence (in negative-sense viral orientation), such that the 5’ coding, random barcode region, and priming sequence were identical to the library strain. A T7 promoter sequence was added after the 5’ NCR for in vitro transcription of the RNA sequence. See https://github.com/dms-vep/flu_h3_hk19_dms/blob/main/library_design/plasmid_maps/rna-spikein-std_template_DNA.gb for full sequence of the linear DNA template. The GFP recipient vector was digested with XbaI alone, then gel and Ampure bead purified. The C-terminal insert sequence was PCR-amplified off of the chimeric library plasmid, using a forward primer that appended a 20-bp overlap with the GFP sequence. PCR conditions were as follows: PCR mix: 21µL H2O, 1.5µL 10µM forward GFP overlap primer (primer_115), 1.5µL 10µM reverse linearizing primer (primer_089), 1µL 10ng/µL template plasmid, and 25µL 2x KOD Hot Start Master Mix. Cycling conditions: (1) 95C/2min (2) 95C/20sec (3) 63C/10sec (4) 70C/10sec (5) Return to Step 2 x19 (6) 70C/2min. Amplified insert was gel purified.

We then appended random 16-nucleotide barcodes to the amplified insert, followed by a constant overlap sequence for cloning into the GFP recipient vector, using a PCR with the following conditions: PCR mix: 10.66µL H2O, 0.9µL 10µM forward linearizing primer (primer_088), 0.9µL 10µM reverse barcoding primer (primer_090), 2.54µL 11.8ng/µL insert template, and 15µL 2x KOD Hot Start Master Mix. Cycling conditions: (1) 95C/2min (2) 95C/20sec (3) 70C/1sec (4) 50C/30sec, cooling at 0.5C/sec (5) 70C/20sec (6) Return to Step 2 x9 (7) 70C/2min.

The barcoded insert was gel purified, then cloned into the XbaI-digested GFP recipient vector, using a 3:1 insert to vector ratio in a 1 hour HiFi assembly reaction. NEB 5α High Efficiency Cells were transformed with 2µL of the HiFi assembly, and plated on LB+ampicillin for 18 hours of outgrowth. To generate a diverse pool of barcodes, 20 individual colonies were grown in liquid LB+ampicillin at 37°C for 18 hours, and plasmids were purified using QIAprep Spin Miniprep Kit. Sequence was confirmed by Sanger sequencing. The 16 plasmids with no SNPs were pooled at equal concentrations.

From this randomly-barcoded pool, we amplified the linear DNA template from the U12 to U13 region, and added a T7 promoter sequence in the reverse orientation after U13 (5’ terminus of the linear template, negative-sense viral orientation). PCR conditions were as follows: PCR mix: 11.2µL H2O, 0.9µL 10µM forward U12-annealing primer (primer_113), 0.9µL 10µM reverse T7-appending primer (primer_116), 2µL 10ng/µL template plasmid, and 15µL 2x KOD Hot Start Master Mix. Cycling conditions: (1) 95C/2min (2) 95C/20sec (3) 62C/10sec (4) 70C/40sec (5) Return to Step 2 x19 (6) 70C/2min. We ran two replicate 30uL PCR reactions, and gel purified both reactions using the same column to maximize concentration of the linear DNA template.

The RNA spike-in standard was then generated via *in vitro* transcription of the linear DNA template with T7 promoter, using the T7 RiboMAX Express Large Scale RNA Production System (Promega, Cat. No. P1320) with 1µg DNA template input. Reaction was incubated at 37°C for 45min. To remove DNA template after transcription, we added 1µL of RQ1 RNase-Free DNase and incubated for an additional 15 min at 37°C. RNA was then purified using the Monarch RNA Cleanup Kit (NEB, Cat. No. T2040).

Final RNA yield was at a concentration of 2.24µg/µL. This was diluted to 0.025ng/µL, aliquoted, and stored at -80°C. Barcodes were amplified and sequenced on a NextSeq 2000 alongside library selection samples, as described in the section ‘Barcode amplification from infected cells for Illumina sequencing’. The final pool of RNA spike-in neutralization standard contained 125 unique barcodes.

### Serum preparation

All sera were heat-inactivated and treated with receptor-destroying enzyme (RDE) before use. RDE-treatment removes residual sialic acids in human serum samples that can bind to the virus libraries. We mixed serum with resuspended, lyophilized RDE II (VWR, Cat. No. 370013) at a ratio of 1:3, and incubated in heatblocks at 37°C for 2.5hr. We then heat-inactivated the RDE and serum by incubating at 55°C for 30min, aliquoted, and stored at -80°C. Note that all sera used in this study are at a starting dilution of 1:4 due to RDE treatment, which is accounted for in reported concentrations.

### GFP neutralization assays for serum screening and validation

Neutralization assays were performed using influenza viruses carrying GFP in the PB1 segment. These GFP viruses were generated by reverse genetics using the following plasmids: wildtype WSN-flanked A/Hong Kong/45/2019 HA (the library background strain), pHH-PB1flank-eGFP (which encodes GFP on the PB1 segment; (Bloom et al., 2010)), and the pHW18* series of plasmids (Hoffmann et al., 2000) for all viral genes except HA and PB1. The same approach was used to generate GFP virus for validation neutralization assays, with the mutation of interest introduced into the WSN-flanked A/Hong Kong/45/2019 HA plasmid.

We added 4x10^5^ 293T-CMV-PB1 (Bloom et al., 2010) and 5x10^4^ MDCK-SIAT1-CMV-PB1-TMPRSS2 (Lee et al., 2018) cells in D10 media to each well of a 6-well plate. Cells that constitutively express PB1 are necessary because the PB1 segment is replaced by GFP in these viruses. 18 hours after plating, wells were transfected with 220ng of each reverse genetics plasmid and the pHAGE2-EF1aInt-TMPRSS2-IRES-mCherry plasmid, using BioT transfection reagent. At 20 hours post-transfection, we changed the media in each well to 2mL NAM (Neutralization Assay Media, consisting of Medium-199 supplemented with 0.01% heat-inactivated FBS, 0.3% BSA, 100 U/mL penicillin, 100 μg/mL streptomycin, 100 μg/mL calcium chloride, and 25mM HEPES). This media is used because it has low background fluorescence at 488 nm. At approximately 40 hours post-transfection, the transfection supernatants were harvested, clarified by centrifugation at 2,000x*g* for 5min, aliquoted, and frozen at -80°C.

To titer these viruses, we infected 1x10^5^ MDCK-SIAT1-CMV-PB1-TMPRSS2 in a 12-well plate with different dilutions of virus, chose wells that showed 1-10% GFP positivity at 18 hours post-infection, measured the fraction of GFP-positive cells by flow cytometry, and used this fraction to calculate the titer of infectious particles per µL.

For accurate measurement of serum neutralization curves, the correlation between number of infectious particles and change in GFP fluorescence should be linear. Adding too much or too little virus per well will result in inaccurate readings. It’s therefore important to generate an MOI curve for each GFP virus, where MDCK-SIAT1-CMV-PB1-TMPRSS2 cells in a 96-well plate are infected with 2-fold dilutions of virus, and GFP signal is measured 18 hours post-infection. The virus concentration used for neutralization assays should correspond to the highest MOI in the linear range of MOI:signal.

Neutralization assays were performed as described previously (Doud et al., 2018; Hooper and Bloom, 2013) (see also https://github.com/jbloomlab/flu_PB1flank-GFP_neut_assay for detailed protocol). Serum was diluted 2-fold in NAM, for final volumes of 40µL per well of a 96-well plate. These dilutions were 2x more concentrated than the target dilutions, to account for the addition of 40µL of virus. Note that calculation of neutralization curves assumes that antibody is in excess of virus, and if this is not the case, the measured IC50 will be artificially high. For some highly potent sera (IC50=1x10^-4^ dilution or lower), serum dilution volumes were therefore increased to 100µL to avoid saturation of serum antibodies. The serum-virus mix was incubated for 1.5 hours at 37°C before adding 4x10^4^ MDCK-SIAT1-CMV-PB1-TMPRSS2 cells per well. When screening for neutralization activity against the wildtype library strain, only one replicate was run per serum. Sera selected for escape mapping were then re-run with two replicates per serum to generate more accurate curves. All validation neutralization assays were run with two replicates per serum and virus, and curves represent the mean and standard error of these replicates.

GFP signal was measured 18 hours post-infection. To calculate fraction infectivity, we subtracted the average background signal from uninfected cells from each condition, then divided the signal from each serum dilution by the average signal from no-serum control infections. Fraction infectivities were then used to fit Hill-like neutralization curves using the neutcurve package (https://jbloomlab.github.io/neutcurve/), version 0.5.7. See https://github.com/dms-vep/flu_h3_hk19_dms/tree/main/neut_assays for detailed information on curve fitting for all neutralization assays in this paper.

### Serum selection of library escape variants

Selections were performed at three concentrations for each serum, targeting a range of concentrations from serum IC90 to IC99.99. Using multiple potent serum concentrations for selections generates more accurate measurements of enriched escape variants, and improves fitting with our biophysical escape models (Yu et al., 2022).

For each experiment, we plated MDCK-SIAT1 cells in IGM 4 hours before infection. Because large differences in MOI impact the amount of viral mRNA transcribed by each cell (Bacsik et al., 2023), we plated 1.5x10^5^ cells per well for each selection condition, and 1x10^6^ cells per well for the no-antibody controls, to roughly account for the expected difference in infectious particles. Selections used 12-well plates while no-antibody controls used 6-well plates, to keep cell density relatively consistent. We made a master mix of library virus with WSN-H6 HA neutralization standard added at 0.1% (libA) or 0.3% (libB) of the total virus titer. These ratios were calibrated in pilot selections such that approximately 1% of counts in the no-antibody control conditions would come from the neutralization standard. Using a master mix of virus is important as it ensures comparable results between the no-antibody and serum selection conditions. 400µL of virus master mix was mixed with 400µL serum (diluted to 2x the targeted concentration, to account for virus volume), or 400µL media for the mock selection control. All dilutions were in IGM. The virus-serum or virus-media mixes were incubated at 37°C for one hour, then added dropwise to each well. RNA was harvested from infected cells at 13 hours post-infection (described below).

In initial pilot experiments, we added 6x10^5^ TCID50 per infection for each library for approximately 20-fold coverage of variant diversity, and ran selections in minimal volumes to conserve serum. We found that these conditions resulted in serum antibody saturation, where there were more virions than antibodies, and results were skewed by measurement of unbound virions. 800µL was the minimal selection volume that avoided saturation effects for potent sera. We also found that the calculated probability of escape for wildtype variants was consistently higher for libB than libA. We therefore reduced the titer of libB added per infection to 3x10^5^ TCID50, which led to consistent results between libraries for wildtype variants. This discrepancy is likely due to inconsistency between TCID50, measured as virus spread in cell culture, and selection experiment results, measured based on cell entry. We therefore recommend using the probability escape for wildtype variants from a pilot experiment, which should reflect targeted ICXX values (e.g. 10% probability escape at serum IC90), to calibrate the amount of virus added per selection.

To minimize bottlenecking of variant diversity during infections, we infected 3 wells with no-antibody controls for each library, prepped the RNA and viral barcodes separately (described below), and pooled the Illumina sequencing reads into a single no-antibody control sample.

### Barcode amplification from infected cells for Illumina sequencing

RNA was harvested from infected cells 13 hours post-infection using the RNeasy Plus Mini Kit (Qiagen, Cat. No. 74134). We made a master mix of RLT lysis buffer with 1% β-mercaptoethanol to inhibit RNases, and 0.14ng/mL RNA spike-in standard (i.e. 2µL of 0.025ng/µL standard for every 350µL in the master mix). This correlates to 0.05ng of standard in 352µL lysis master mix, added to each well. Working with one row of wells at a time, we removed the media, washed with 1mL PBS, and aspirated completely. We then added 352µL of the lysis master mix with RNA standard. Cells were washed from each well and transferred to 1.5mL aliquots, then vortexed for 1min at full speed to homogenize. After lysing and homogenizing cells from all conditions in the experiment, we proceeded with RNA extraction, following the extraction kit protocol. RNA was eluted twice from each column in 30µL RNase-free water to maximize yields. Extraction was typically completed within 1.5 hours from homogenization.

Viral mRNA was transcribed as cDNA, using SuperScript III First Strand Synthesis SuperMix (Invitrogen, Cat. No. 18080400). We normalized amount of input RNA across all samples in the experiment, and input anywhere from 800ng-1µg RNA per sample, depending on the maximum input from the lowest-concentration sample. Reaction was carried out according to manufacturer protocol, using the mRNA-annealing RT primer (primer_110) to amplify off of viral mRNA. We chose to target mRNA as it is more abundant in infected cells than vRNA.

We then used two rounds of PCR to amplify the barcode region and add the necessary Illumina adapters and indices for each sample. The first round of PCR uses a forward primer that anneals to the C-terminus of the HA sequence, and a reverse primer that anneals to the Illumina Truseq Read 1 sequence downstream of the barcode. We use the maximum recommended input of cDNA, 10µL, to minimize bottlenecking of barcode diversity during amplification. Conditions were as follows: PCR mix: 14µL H2O, 0.5µL 5µM Illumina round 1 forward primer (primer_098), 0.5µL 5µM Illumina round 1 reverse primer (primer_099), 10µL cDNA template, 25µL KOD. Cycling conditions: (1) 95C/2min (2) 95C/20sec (3) 70C/1sec (4) 64C/10sec, cooling at 0.5C/sec (5) 70C/20sec (6) Return to Step 2 x19 (7) 70C/2min.

DNA concentration was quantified using a Qubit Fluorometer (ThermoFisher). A second round of PCR was then performed using custom dual indexing primers, which anneal to the Read 1 or Read 2 sites, and append the P5 Illumina adapter and i5 sample index at the Read 1 terminus, or the P7 Illumina adapter and i7 sample index at the Read 2 terminus. Conditions were as follows: PCR mix: 5ng of round 1 PCR product (variable volumes), H2O to 19µL, 3µL 2.5µM unique i5 indexing primer, 3µL 2.5µM unique i7 indexing primer, 25µL KOD. Cycling conditions: (1) 95C/2min (2) 95C/20sec (3) 70C/1sec (4) 55C/20sec (5) 70C/20sec (6) Return to Step 2 x9 (7) 70C/2min.

DNA concentration of each round 2 PCR product was quantified using Qubit. Serum selection samples were pooled at an even ratio, while no-antibody control samples were added at 3x the concentration of each selection sample, to account for greater barcode diversity in the no-antibody controls. Samples were gel purified and Ampure bead cleaned at a 1:2 sample to beads ratio, then sequenced using either P2 or P3 reagent kits on a NextSeq 2000. Analysis is described in the section ‘Illumina barcode sequencing data analysis’.

#### COMPUTATIONAL METHODS

### Computational pipeline overview

For analysis of A/Hong Kong/45/2019 deep mutational scanning data, we used the modular pipeline https://github.com/dms-vep/dms-vep-pipeline (version 2.6.5). The repository https://github.com/dms-vep/flu_h3_hk19_dms contains all analyses performed in this paper, including the main *dms-vep-pipeline* and all scripts, notebooks, and settings necessary to recreate the analysis. Full analysis of A/Perth/16/2009 deep mutational scanning data can be found at https://github.com/jbloomlab/map_flu_serum_Vietnam_H3_Perth2009 and does not use *dms-vep-pipeline* for analysis. However, we have imported the finalized escape scores and validation neutralization data to the main https://github.com/dms-vep/flu_h3_hk19_dms repository, and integrated these results into the pipeline. See https://dms-vep.github.io/flu_h3_hk19_dms/ for HTML rendering of key analyses and interactive plots. These pages are the best way to explore the analyses and interactive plots of the results described in this paper.

### PacBio sequencing data analysis

PacBio CCSs were analyzed using *alignparse* (see https://jbloomlab.github.io/alignparse/ for documentation) (Crawford and Bloom, 2019). We computed the empirical accuracy of the CCSs as the frequency at which CCSs with the same barcode were associated with the same variant sequence (see https://jbloomlab.github.io/alignparse/alignparse.consensus.html#alignparse.consensus.empirical_accuracy for implementation). Empirical accuracies for both libraries were approximately 0.99.

We filtered out any barcodes that had less than two CCSs, to ensure confidence in the consensus sequences. We also filtered out barcodes that had minor fractions of substitutions or indels above 0.2, which would indicate that multiple variants share that barcode. See https://jbloomlab.github.io/alignparse/alignparse.consensus.html#alignparse.consensus.simple_mutconsensus for method for building consensus sequences using these filters. The resulting consensus sequences were saved as barcode / variant lookup tables. This file and the associated analysis notebook, which includes quality control plots, can be found on the main HTML page described in ‘Computational pipeline overview’.

### Illumina barcode sequencing data analysis

We processed the Illumina barcode sequencing using the parser https://jbloomlab.github.io/dms_variants/dms_variants.illuminabarcodeparser.html to determine the counts of each variant in each selection condition. We filtered barcoded mutants that had low counts in the no-antibody control conditions, as these were likely to be randomly bottlenecked during infections and generate noisy escape scores. See https://github.com/dms-vep/flu_h3_hk19_dms/blob/main/config.yaml for the configuration file where these thresholds are specified.

### Modeling the effects of mutations on H3 HA function

We modeled the functional effects of mutations on A/Hong Kong/45/2019 H3 HA as previously described (Dadonaite et al., 2023). We computed functional scores based on the frequency of a given mutation in the plasmid library (where there is no functional pressure), compared to the passaged virus library (where non-functional HA variants have been purged). The functional score for a variant *v* is calculated as *log*_2_ ([*n_post_^v^*/*n^wt^_post_*]/[*n_pre_^v^*/*n^wt^_pre_*]) where *n_post_^v^* is the variant count in the no-antibody mock selection condition, *n_pre_^v^* is the variant count in the plasmid library, and *n^wt^_post_* and *n^wt^_pre_* are the counts of all wildtype barcodes in each condition.

Many variants in the library carry multiple mutations. We therefore use global epistasis models (Otwinowski et al., 2018; Sailer and Harms, 2017) to deconvolve this data and estimate the fitness effects of each individual mutation. Under these models, each mutation has an additive effect on a “latent” (unmeasured) phenotype. These additive effects are then transformed by a nonlinear function to generate the “observed” (measured) phenotype. The nonlinear transformation accounts for epistatic interactions between mutations, and thereby generates more accurate estimates for mutation effects.

See https://dms-vep.github.io/flu_h3_hk19_dms/fit_globalepistasis.html for the notebooks used to fit global epistasis models, https://dms-vep.github.io/flu_h3_hk19_dms/muteffects_latent_heatmap.html for the latent effects of mutations averaged across selections, and https://dms-vep.github.io/flu_h3_hk19_dms/muteffects_observed_heatmap.html for the observed effects of mutations averaged across selections.

### Modeling the effects of mutations on antibody and serum escape

We modeled the effects of mutations on serum escape as described previously in Dadonaite et al. (2023). Briefly, we computed the fraction of non-neutralized variants at each serum selection concentration, normalized by counts from the relevant neutralization standard (either WSN-H6 HA or the RNA spike-in standard). Fraction of non-neutralized variants corresponds to probability of escape, and should fall between 0 (variant is fully neutralized) and 1 (variant fully escapes neutralization). Values greater than 1 are clipped. We then estimated the effects of each individual mutation on escape using the software package *polyclonal*, version 6.2 (see https://jbloomlab.github.io/polyclonal/ for documentation) (Yu et al., 2022). Fits were constrained to a single epitope to simplify analysis, as most sera were highly targeted.

For each serum, the heat maps and line plots are generated using the mutation-level escape scores calculated by *polyclonal*. All sera had measurements from two replicate libraries, and the final reported values are the average (median) across those replicates. Files containing the escape scores from individual replicates, the averaged scores, analysis notebooks with relevant quality control plots, and interactive plots summarizing the final estimates are available on the main HTML page described in ‘Computational pipeline overview’.

Figures in this paper only include mutations seen in at least three distinct variants (averaged across libraries), with a functional effect of at least -1.38.

### Estimating escape scores for circulating H3N2 strains

The GitHub repository at https://github.com/matsengrp/seasonal-flu-dmsa has the code we used to perform the analyses in this section, as well as files with key results. We analyzed a set of 1,478 circulating H3N2 strains obtained from a publicly available Nextstrain (Hadfield et al., 2018) phylogenetic tree that samples strains from the past 12 years (https://nextstrain.org/flu/seasonal/h3n2/ha/12y). This tree is regularly updated. For this paper, we analyzed the strains from the tree that was available on October 19th, 2023. For each strain, we obtained the corresponding HA gene nucleotide sequence and associated metadata from the GISAID database (https://gisaid.org/) (Shu and McCauley, 2017). The above repository provides the GISAID accession number for each strain’s HA sequence and acknowledges contributing labs.

To estimate an escape score for each strain relative to each serum sample, we ran the above sequences and metadata, including the sequence and metadata for the HA sequence of the A/Hong Kong/45/2019 strain used in the DMS experiment, through a version of the core *seasonal-flu* Nextstrain pipeline (https://github.com/nextstrain/seasonal-flu/) that we modified to compute escape scores. This pipeline generates a multiple-sequence alignment of translated HA protein sequences. For each strain in the alignment, it then makes a list of all amino-acid mutations that are needed to convert the A/Hong Kong/45/2019 sequence to the sequence of the given strain. Then, to compute that strain’s escape score for a given serum sample, it simply sums the serum-specific DMS-measured escape scores of all mutations in the above list, such that escape scores for the A/Hong Kong/45/2019 DMS strain always equals zero. We only summed escape scores for mutations that were seen in both libraries and in at least three distinct variants (averaged across libraries) in the DMS data for that serum sample. If a strain’s list of mutations included one or more mutations that were not measured by DMS or did not meet the above criteria, we masked that strain in the analysis, unless the mutations in question occurred at sites that we deemed unlikely to affect escape (see the repository for more detail). After the masking step, this process yielded a total of 1,233-1,238 strains with estimated escape scores for each serum sample.

Figure 6B shows the distribution of strain-specific escape scores as a function of strain collection date, with scores averaged across all sera in a given cohort. For each cohort, we fit a linear-regression model that predicts a strain’s average escape score as a function of its collection date and reported the inferred slope of the resulting best-fit line in the figure caption. Next, we used a randomization approach to determine if these slopes were significantly different between cohorts. In each of 1,000 independent trials, we randomized which serum sample was assigned to which cohort, recomputed average strain-specific escape scores using the randomized cohort assignments, and then fit a linear-regression model to the resulting data, fitting one model per cohort as above. We then compared the slopes obtained by randomization to the slopes observed in the non-randomized data. In Figure 6B, the slope associated with teenagers is 0.1 units greater than the slope associated with adults (=0.14-0.04). Out of the 1,000 randomization trials, only four trials resulted in a difference in slope (=teenagers-adults) greater than or equal to 0.1, indicating that the slope for teenagers is significantly larger than the slope for adults in the non-randomized data (p=0.004). The randomization test also indicated that the slope for teenagers is significantly larger than the slope for children (p=0.022), though the slope for children was not significantly larger than the slope for adults (p=0.323).

---

**Figure S1.**
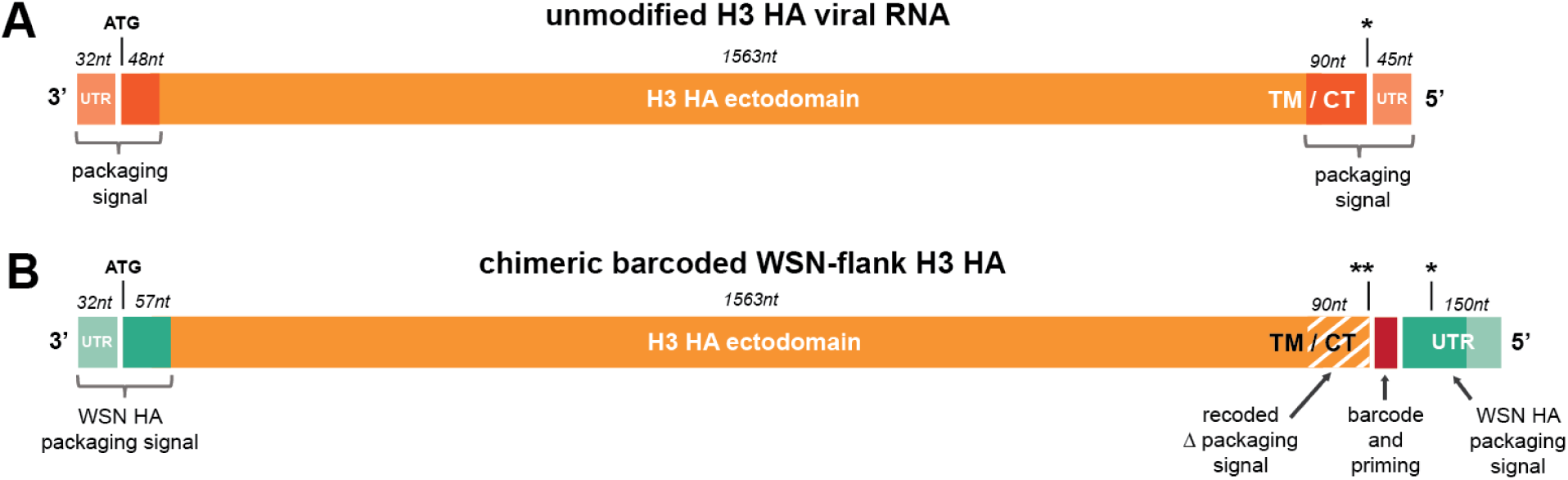
Design of chimeric barcoded WSN-flank H3 HA construct. All segments are shown in the reverse orientation of the negative-sense viral genome, with the 3’ to 5’ labels indicating ends in the negative-sense viral genome. Stop codons are denoted as asterisks. (A) Schematic of normal unmodified H3 HA. Influenza is multi-segmented, and proper packaging of vRNA into the virion relies on segment-specific RNA sequences called “packaging signals.” These span both coding and noncoding regions at the 3’ and 5’ ends of the vRNA segment (Li et al., 2021). Packaging signals, untranslated regions (UTRs), ectodomain, transmembrane domain (TM), and cytoplasmic tail (CT) are labeled for a normal H3 HA. Corresponding nucleotide length is labeled based on sequence of A/Hong Kong/45/2019. (B) Schematic of chimeric WSN-flank H3 HA used for barcoding. To insert a barcode without disrupting vRNA packaging, we duplicate the full 5’ packaging signal (including sequence from the coding region), and place it after the stop codons of the HA gene. The second stop codon was introduced to minimize polymerase read-through. We then insert a 16nt barcode and constant priming sequence after the stop codon and before the intact packaging signal. The native packaging sequence in the 5’ coding region is synonymously recoded to avoid interference between duplicated sequences. Homology between the native and duplicated packaging signal could lead to barcode loss; we therefore included a stop codon in the duplicated packaging signal that will be in-frame if it replaces the native packaging signal. This ensures that loss of the barcode will generate a truncated, non-functional HA protein. The 3’ and duplicated 5’ packaging signals are taken from the lab-adapted A/WSN/1933 HA, as the library is grown using the other seven gene segments from this strain, and consistency between packaging signals helps achieve higher titers. See https://github.com/dms-vep/flu_h3_hk19_dms/tree/main/library_design/plasmid_maps for an annotated plasmid map of WSN-flank H3 HA.

**Figure S2.**
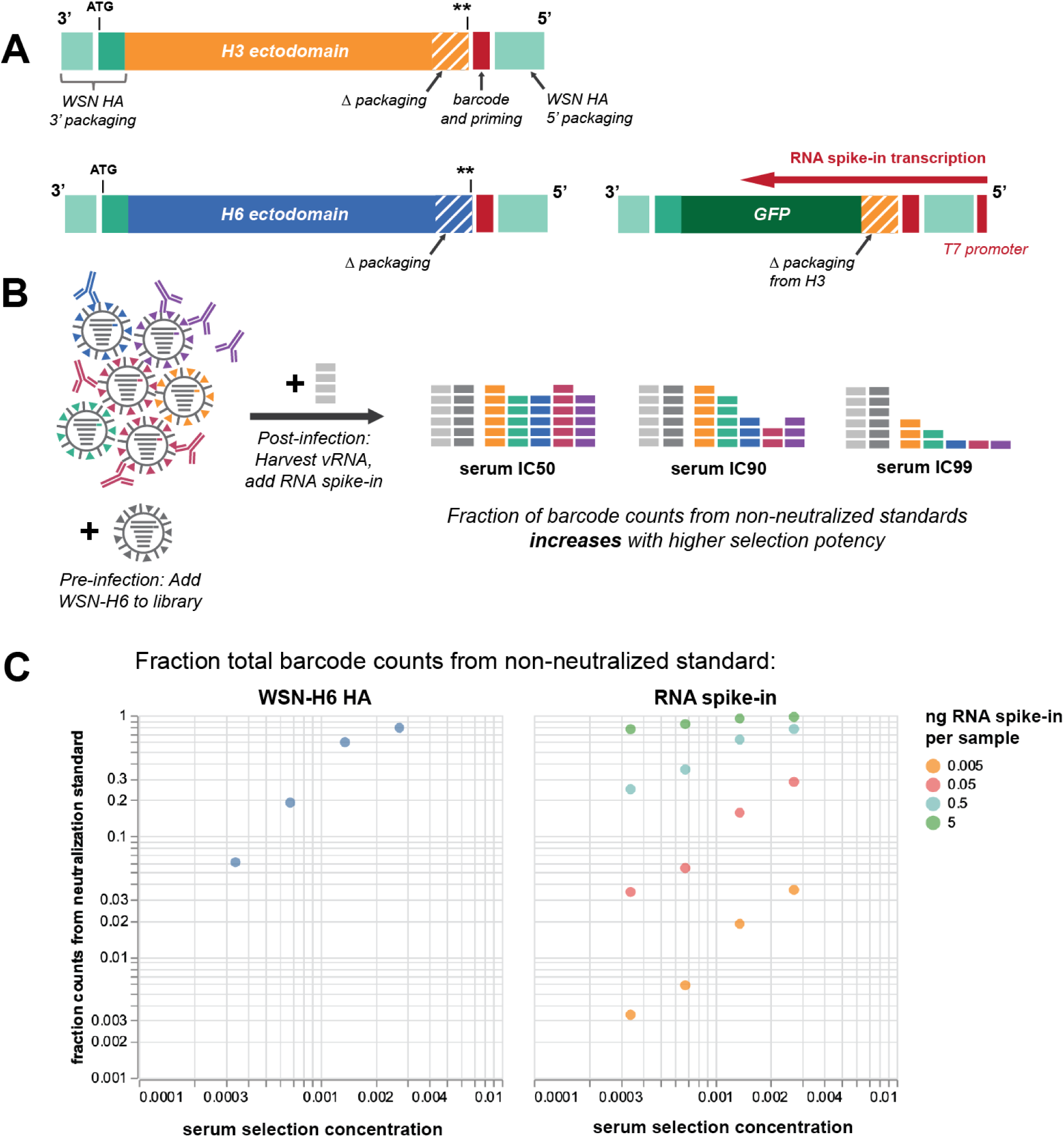
Incorporation of non-neutralized standards to generate quantitative escape measurements. (A) Design of WSN-H6 HA and RNA spike-in neutralization standards. The WSN-H6 HA standard is identical to the H3 HA library construct, but uses the ectodomain and 5’ region from A/Turkey/Mass/1975, a low-pathogenicity avian HA strain (GenBank Accession AB296072.1) (Sandbulte et al., 2009). The RNA spike-in is generated by *in vitro* transcription of a construct where the H3 HA ectodomain is replaced by GFP. See https://github.com/dms-vep/flu_h3_hk19_dms/tree/main/library_design/plasmid_maps for annotated plasmid maps of both standards. (B) WSN-H6 HA is spiked into the H3 HA library before incubating with serum and infecting cells. The RNA spike-in standard is added when harvesting cellular RNA at 13 hours post-infection. As neither standard is neutralized by human sera, the number of barcodes from the standards remains constant, while H3 HA barcodes decrease at higher concentrations of neutralizing antibodies. The overall fraction of standard barcodes should therefore increase at a constant rate with increasing serum potency. (C) At a range of serum concentrations used for selections, the fraction of counts from the WSN-H6 HA neutralization standard increases in a consistent manner as the H3 HA variant library is more potently neutralized. Spiking in 0.05ng of RNA standard per sample achieves the same effect.

**Figure S3.**
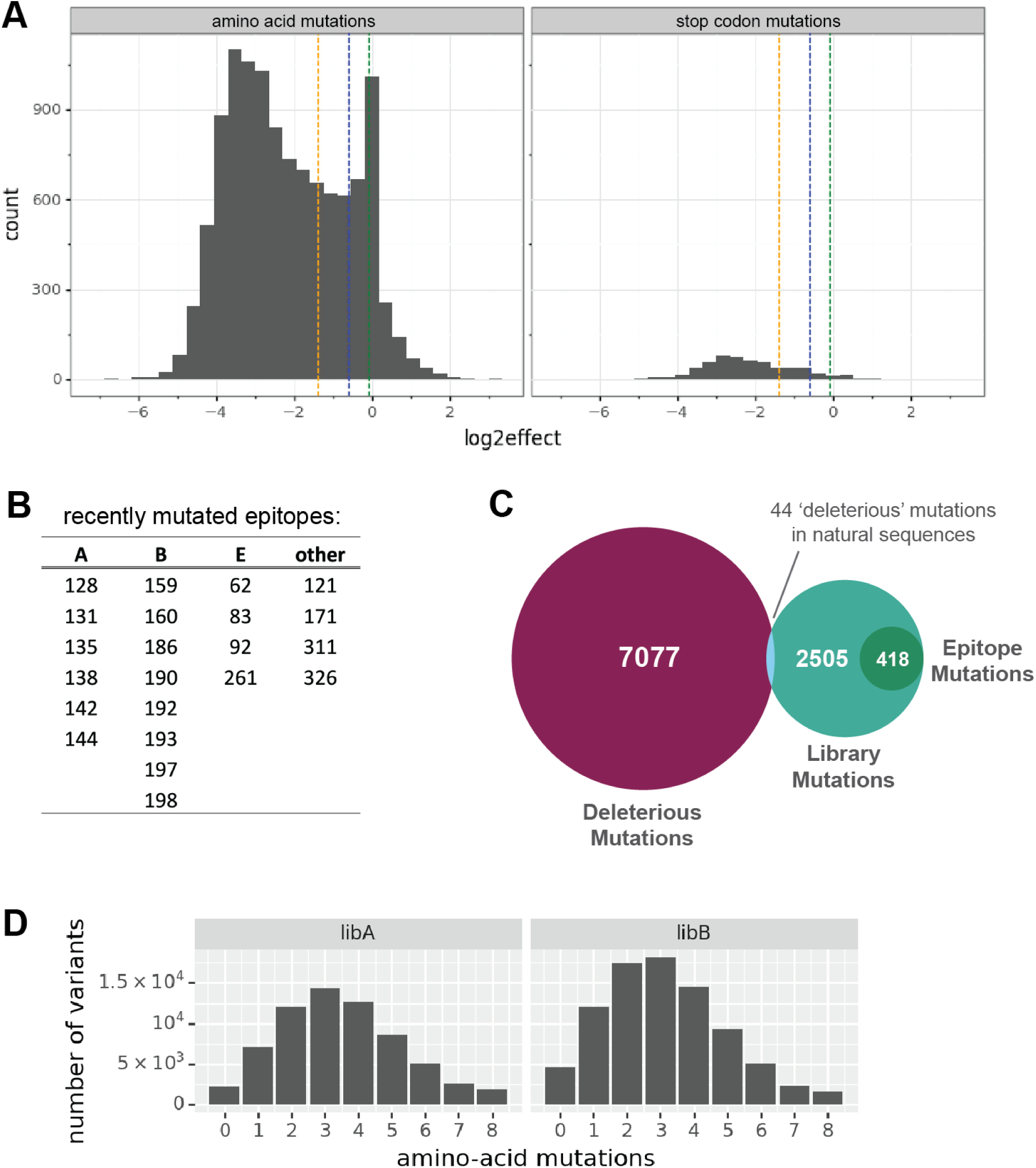
Composition of H3 HA libraries. (A) Functional effects of all single amino acid mutations to A/Perth/16/2009 H3 HA, calculated by Lee, et al. (2018). Vertical lines delimit 75%, 90%, and 95% of the most deleterious stop codons. These quantiles are transposed onto all amino acid mutations, and 7,077 mutants corresponding to the 75th quantile of stop codons were excluded from the current library. (B) List of epitope sites, defined as sites that have been mutated in at least one major H3 HA clade between 2015 and 2021. These sites are randomly mutagenized in the library at a higher rate than the general non-deleterious mutations. (C) Visualization of types of mutations included in the library. In total, 7,077 mutations in the H3 HA ectodomain region were identified as functionally deleterious. 44 of these excluded mutants were identified in H3 HA strains circulating between 1968 and 2021, and added back into the library. This left 2,505 mutations to target in the library. 418 mutations at epitope sites were introduced at a higher frequency than general library mutations. (D) Distribution of mutations per variant in two fully independent plasmid libraries.

**Figure S4.**
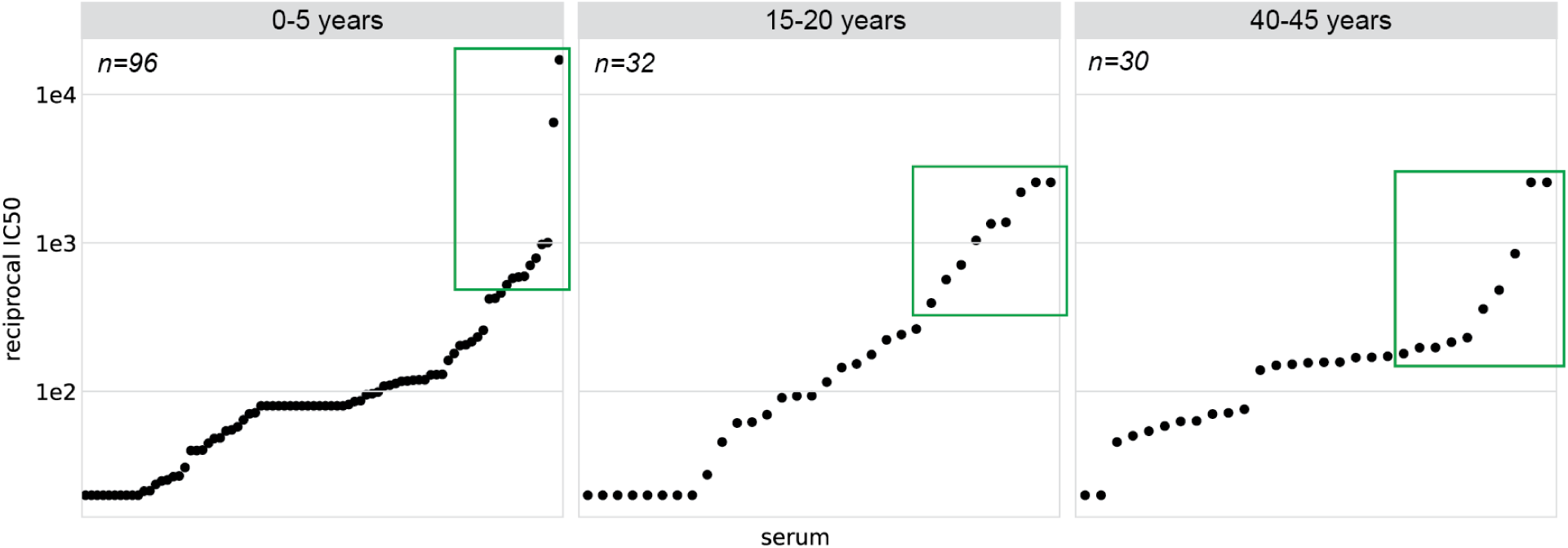
Serum neutralizing titers against the A/Hong Kong/45/2019 HA barcoded library strain. Neutralization assays were performed using influenza carrying GFP in the PB1 segment, as described previously (Doud et al., 2018; Hooper and Bloom, 2013; see https://github.com/jbloomlab/flu_PB1flank-GFP_neut_assay for detailed protocol). Only a single replicate was run for each serum. IC50 values were calculated by fitting Hill-like curves using the neutcurve Python package (https://jbloomlab.github.io/neutcurve/). The ten sera from each age group with the highest neutralization activity against the parental library strain, boxed in green, were selected for serum escape mapping.

**Figure S5.**
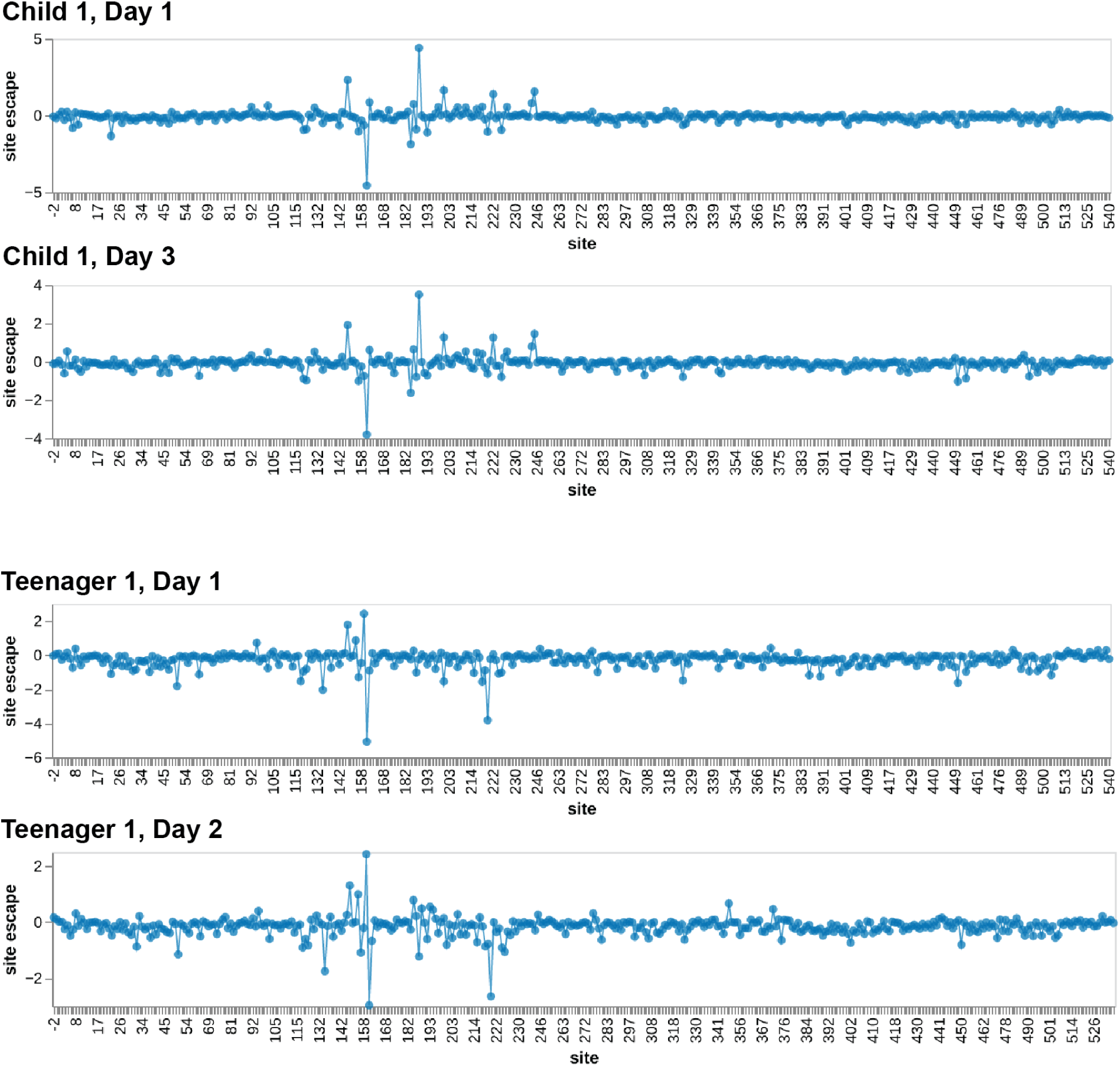
Escape maps for sera collected on different days from the same individual. One set of repeated samples comes from a child (sample 2388), and the other from a teenager (sample 3862). Line plots show summed escape scores of each sampled mutation at that site.

**Figure S6.**
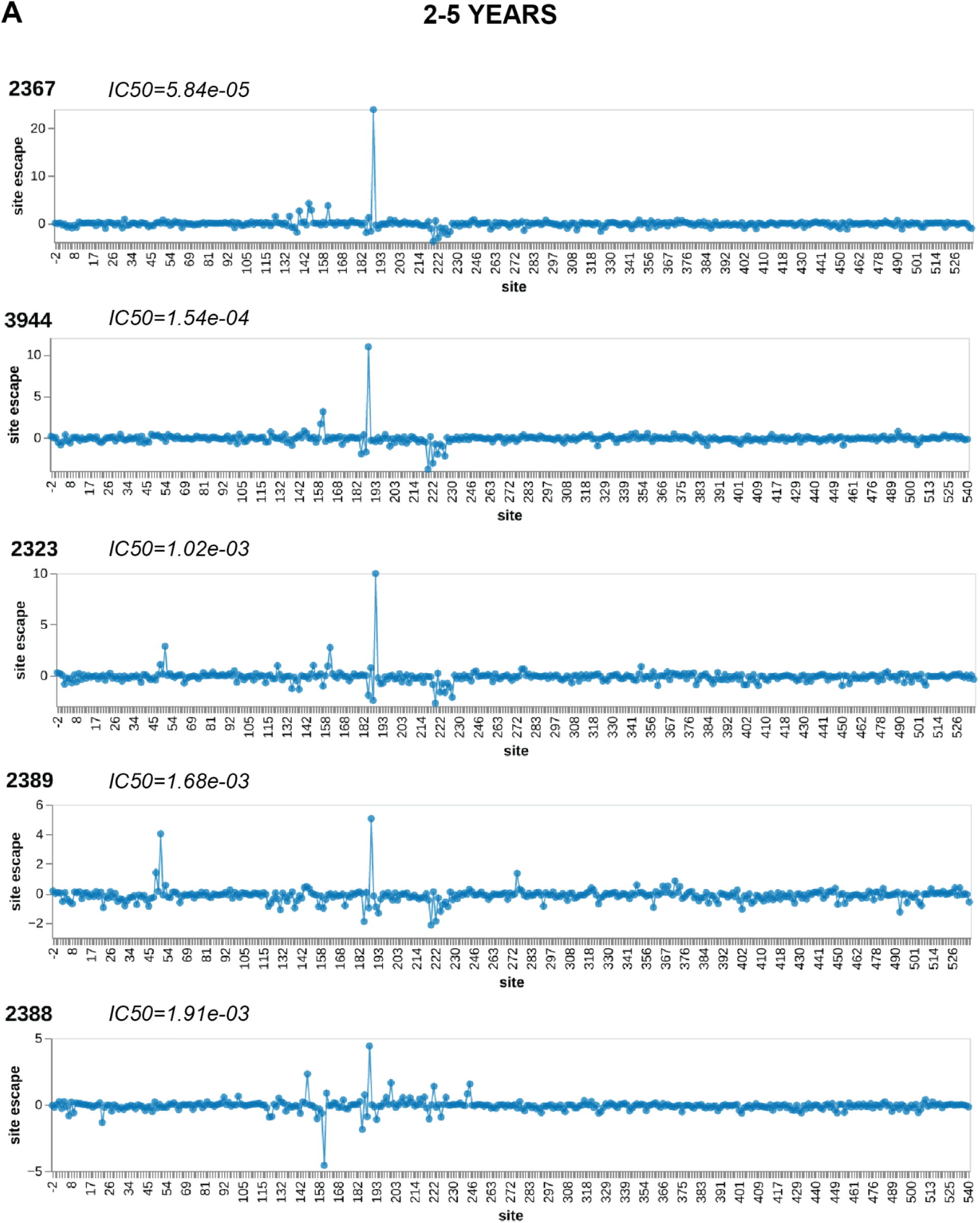
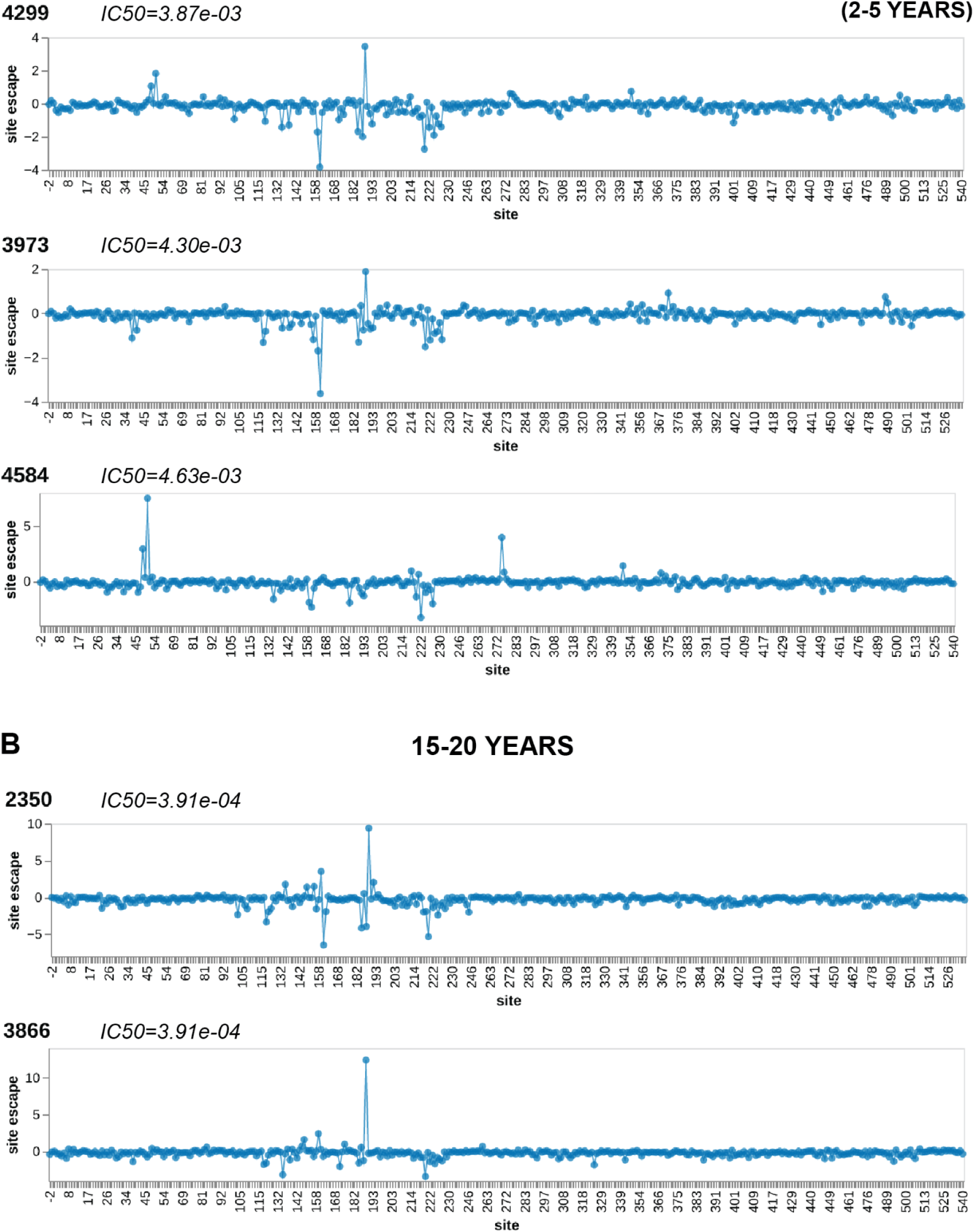
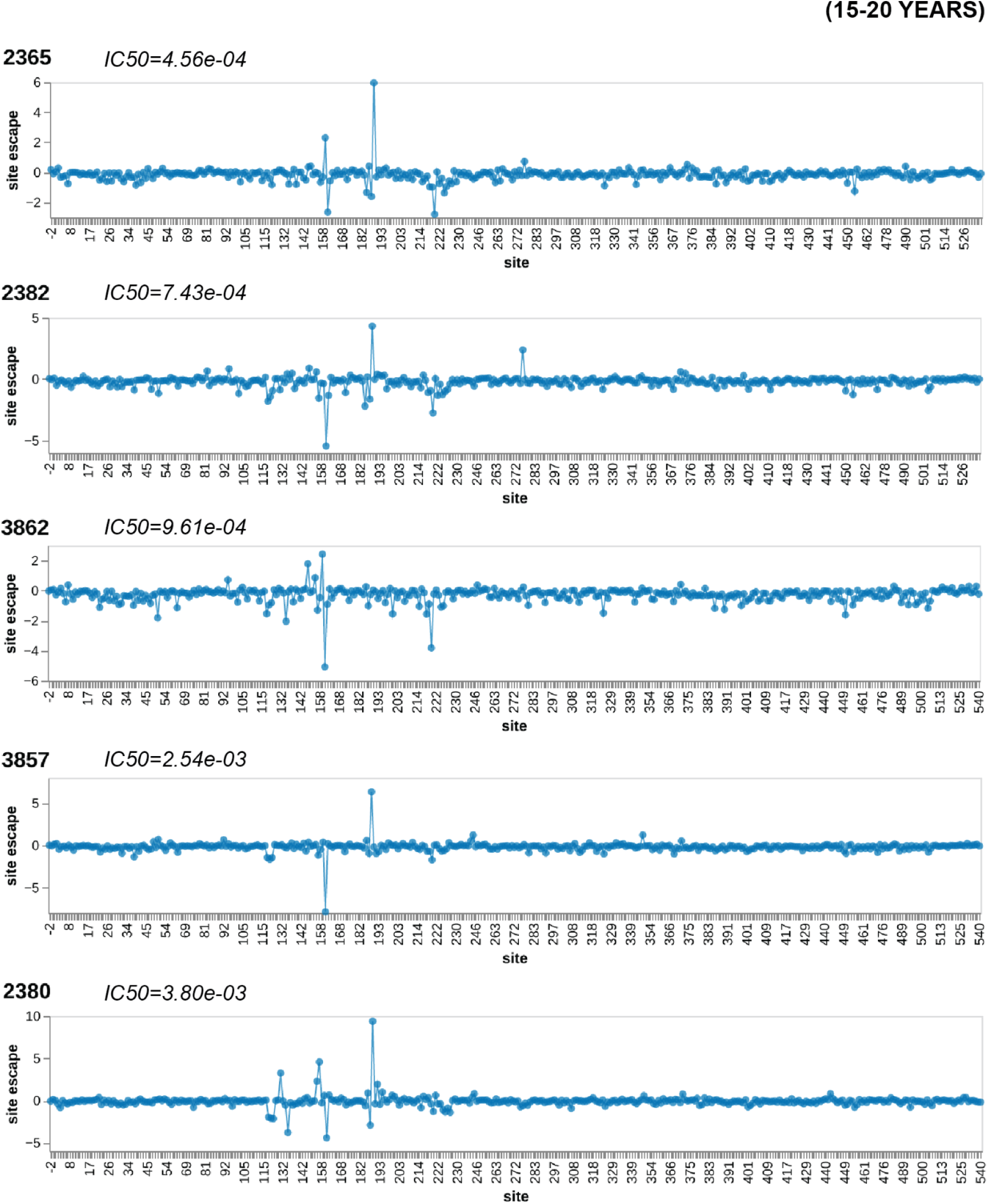
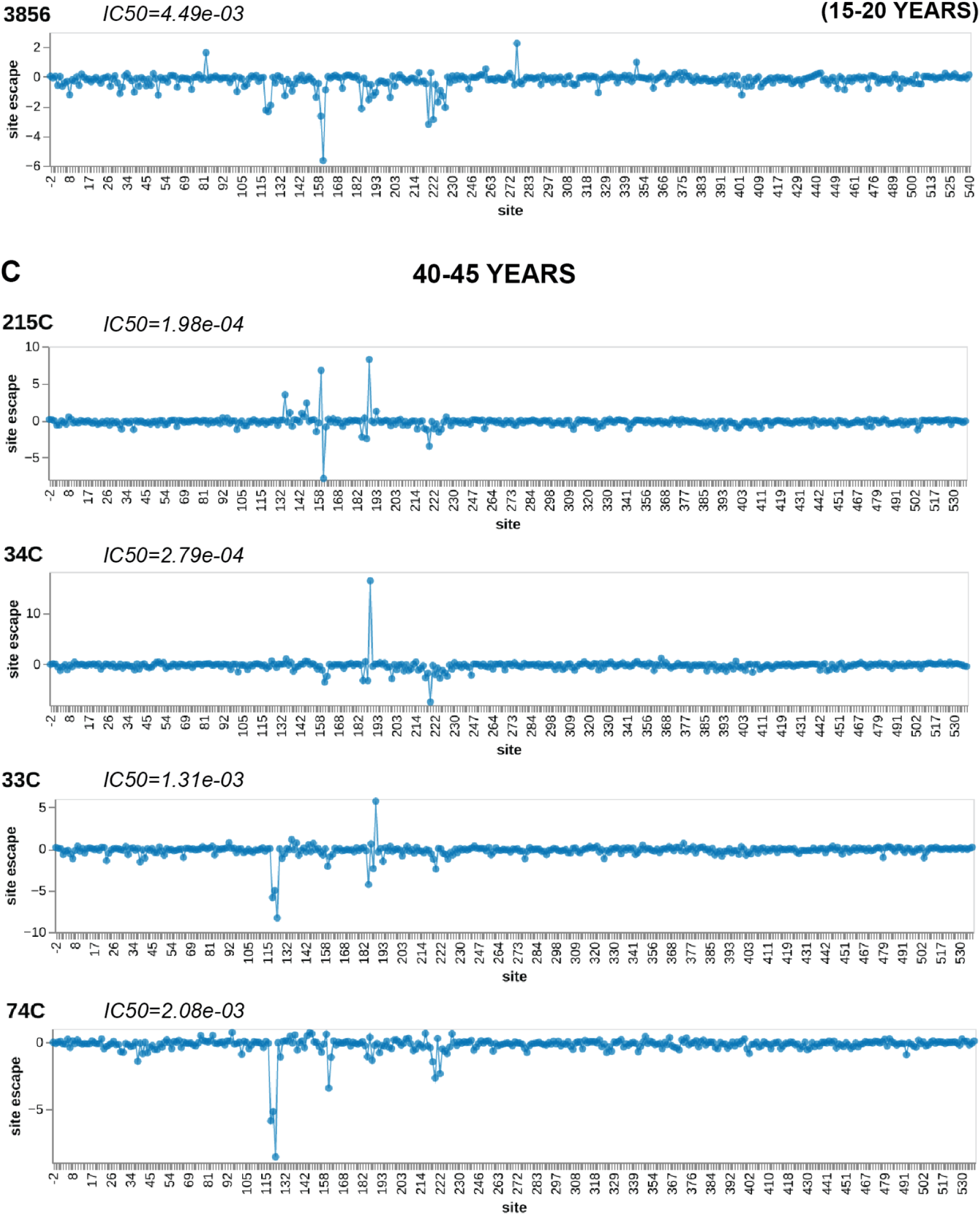
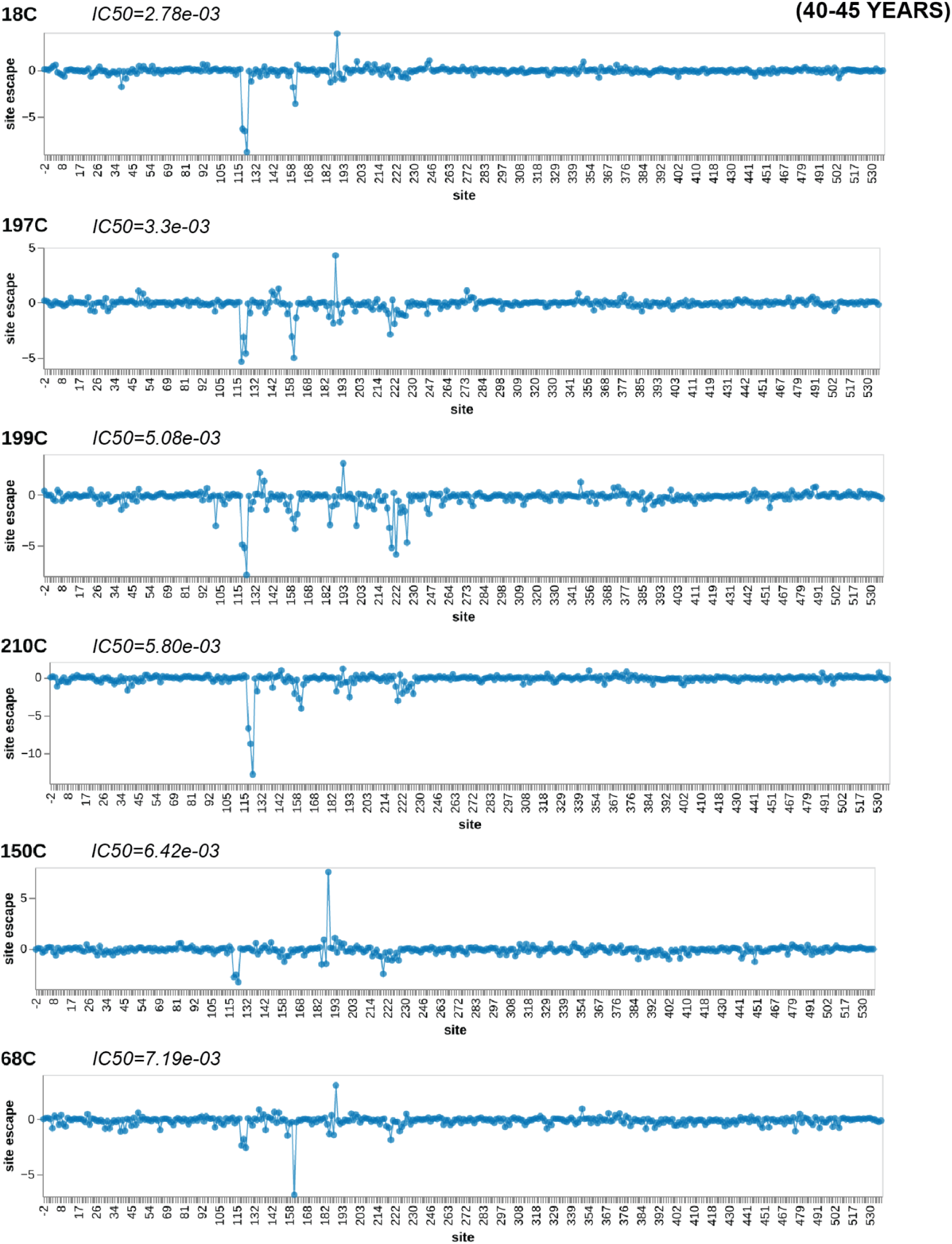
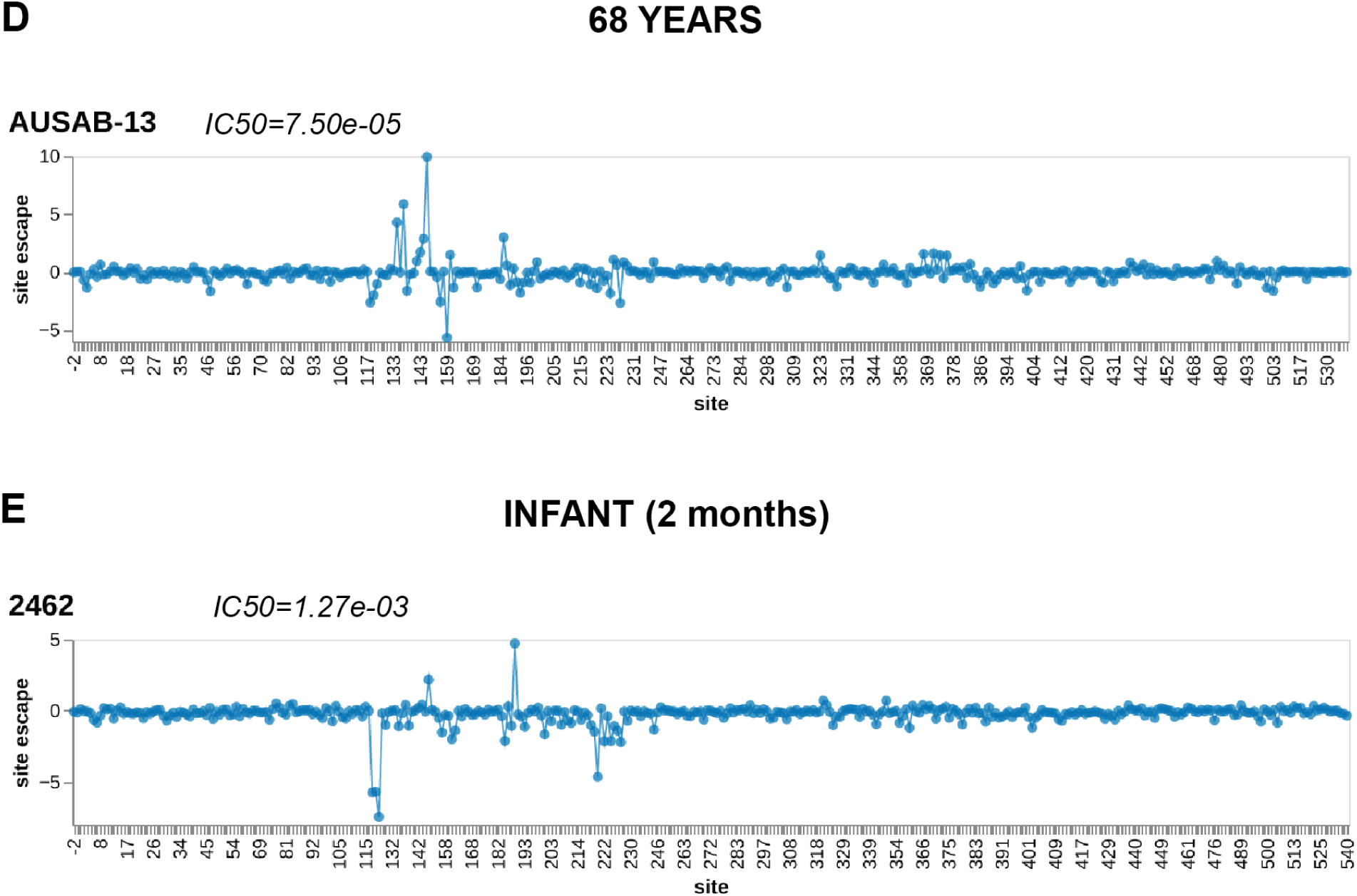
Individual escape maps for each serum in the 2020 cohort, analyzed against the A/Hong Kong/45/2019 library. Line plots show summed escape scores of each sampled mutation at that site. Sera are ordered by decreasing potency within each age group, with IC50s labeled for each serum. (A) Child sera, (B) teenage sera, (C) adult sera, (D) elderly serum, and (E) infant serum.

**Figure S7.**
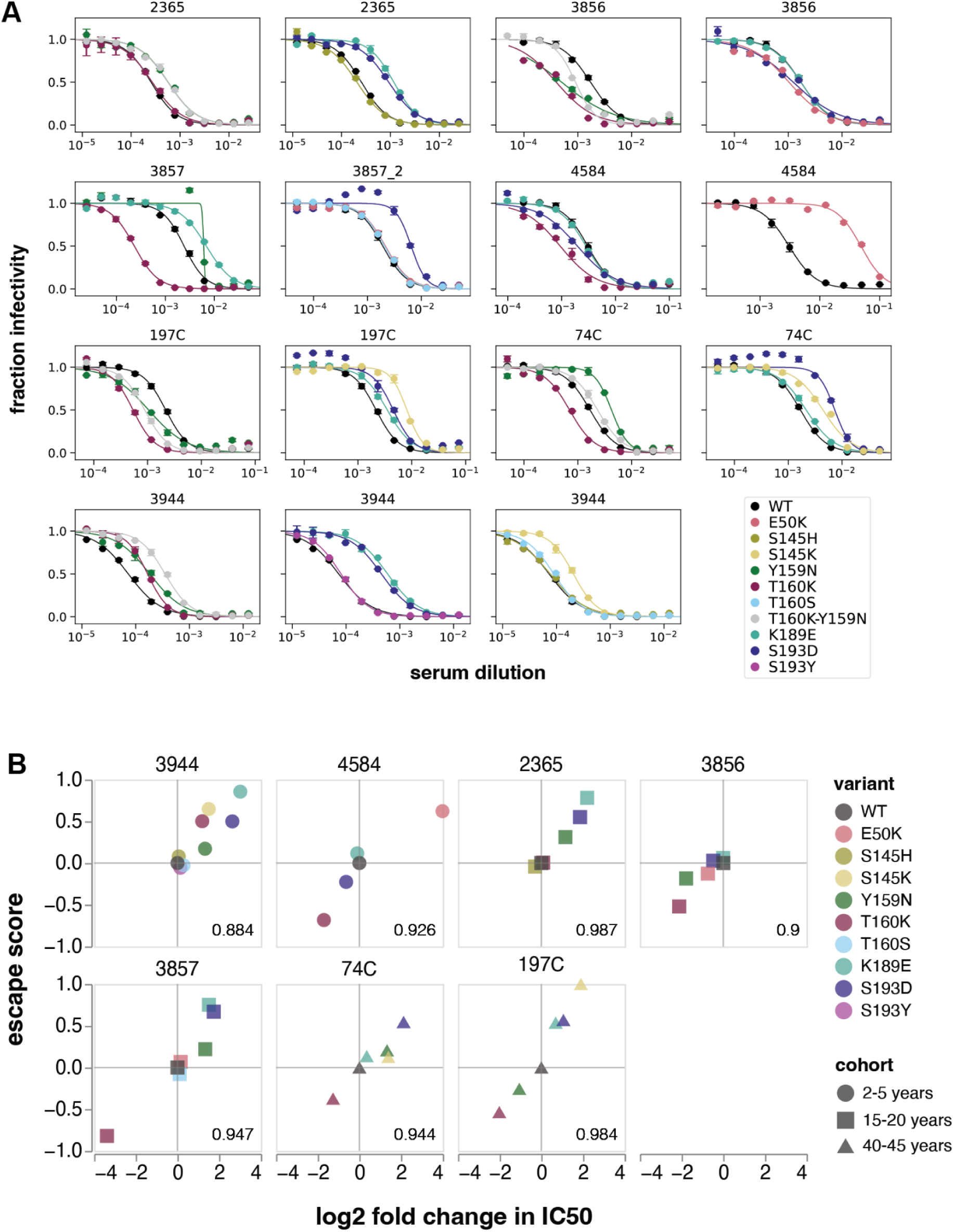
Neutralization assay results for validation of deep mutational scanning measurements against the A/Hong Kong/45/2019 library. (A) Neutralization curves for unmutated A/Hong Kong/45/2019 H3 HA (‘WT’) and selected mutants against seven representative sera from different age cohorts. (B) Correlation between escape score and the log2 fold change in serum IC50 for each variant, plotted for each serum independently. Pearson R correlation is noted on each plot. The IC50 for each variant was compared to the IC50 for the wildtype strain run in the same experiment, to control for potential variation in serum concentration and cell number between independent experiments.

**Figure S8.**
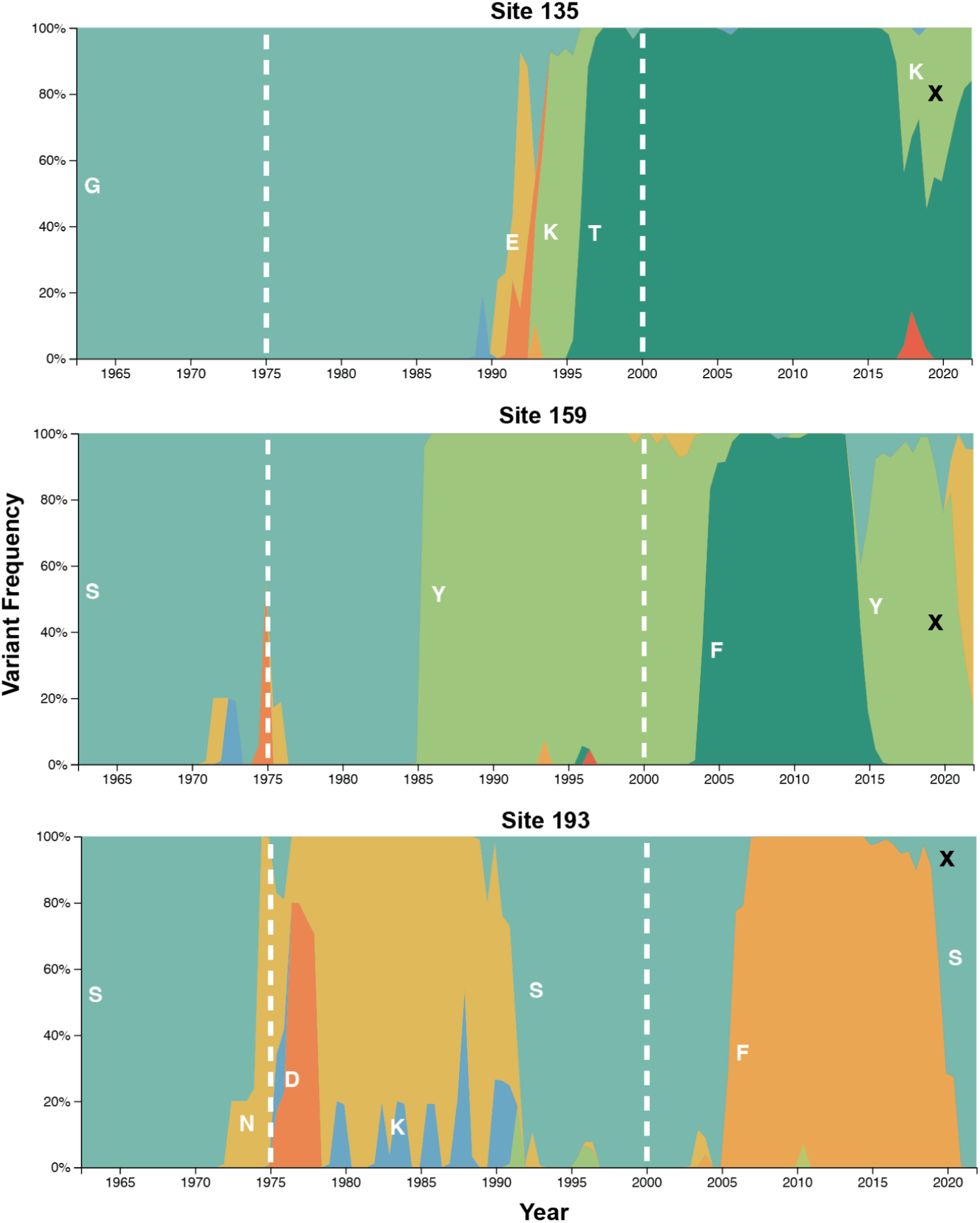
Global frequency of variants at sites 135, 159, and 193. Dashed lines indicate the earliest birthdate of individuals in the adult cohort (1975) and teenage cohort (2000). Black ‘X’ indicates the amino acid identity of the wildtype library strain, A/Hong Kong/45/2019. Frequency plot adapted from the Nextstrain real-time pathogen evolution website (Hadfield et al., 2018; Neher and Bedford, 2015).

**Figure S9.**
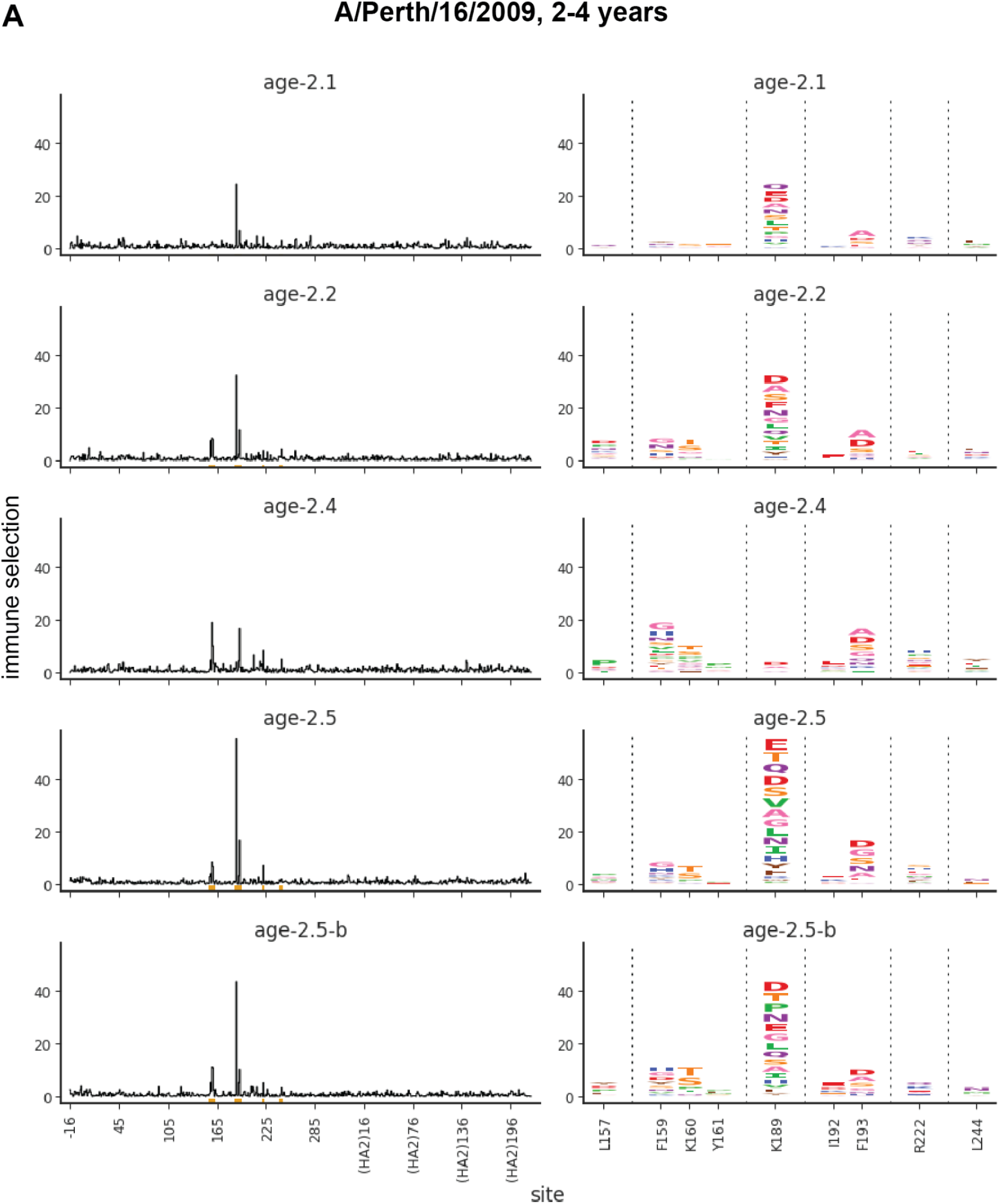
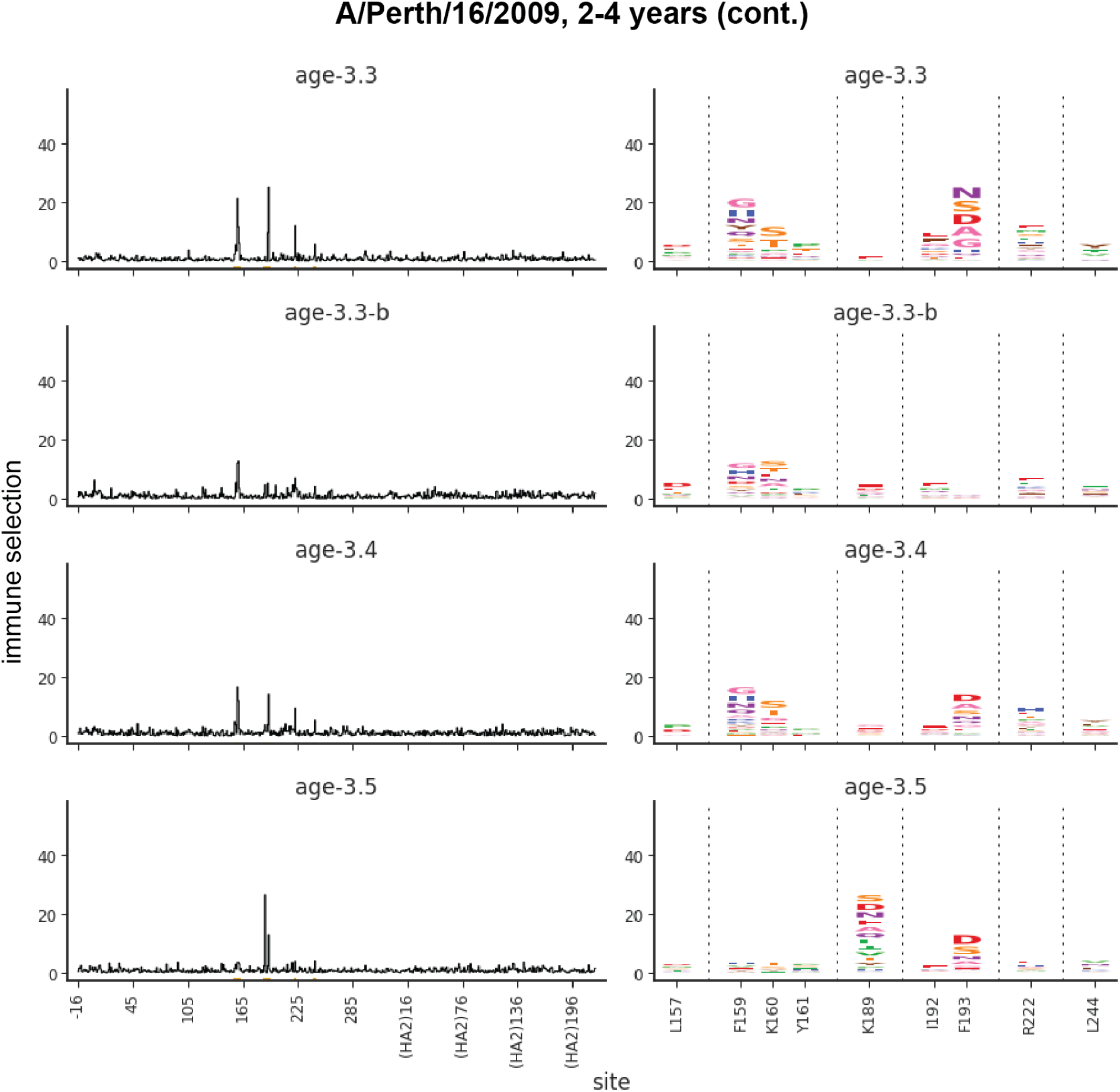
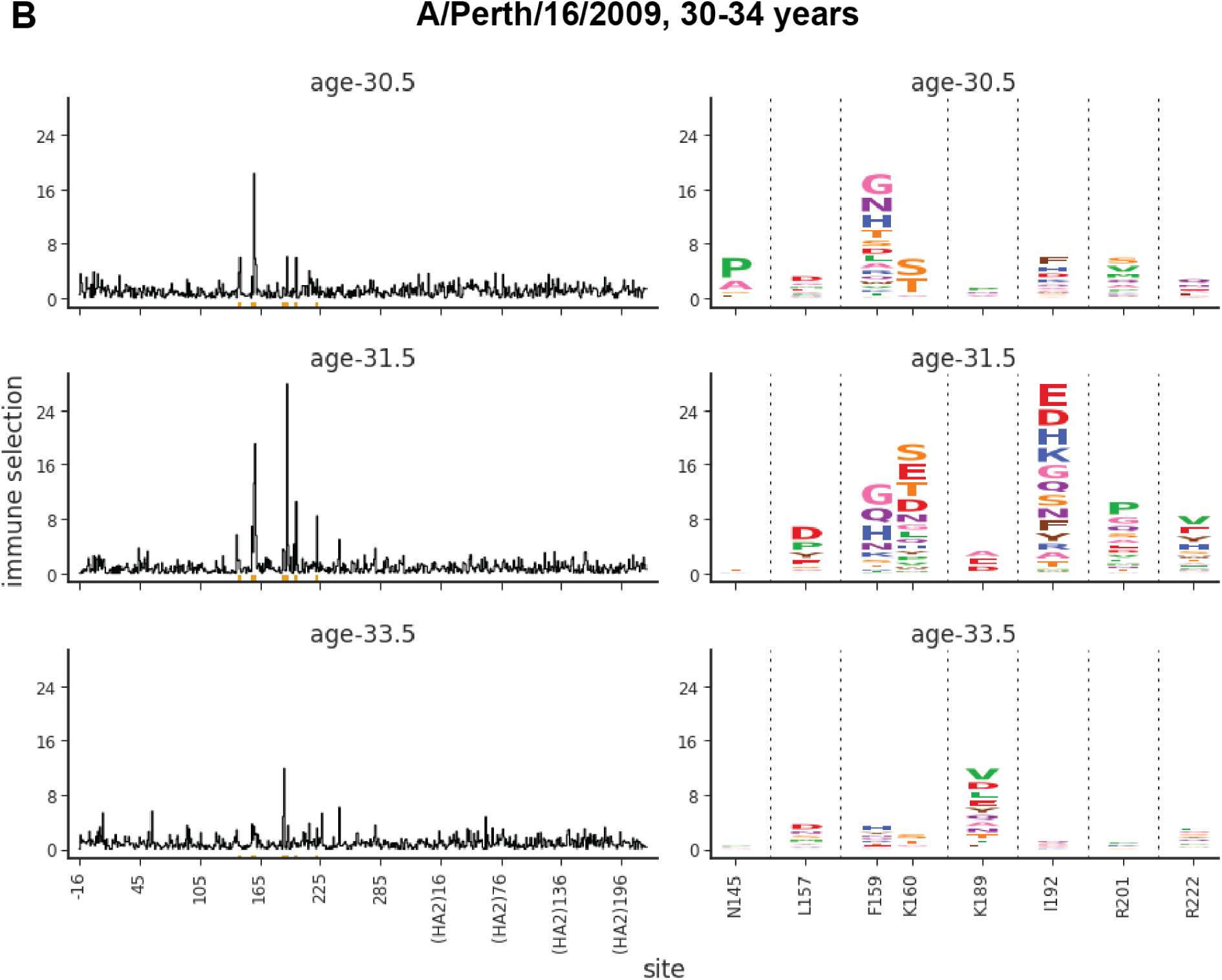
Individual escape maps for each serum in the 2010-2011 cohort, analyzed against the A/Perth/16/2009 library. Line plots show summed escape scores of each sampled mutation at that site. Logo plots show mutation-level escape at key sites, where the height of each letter corresponds to the escape score for that amino acid substitution. (A) Child sera, (B) adult sera.

**Figure S10.**
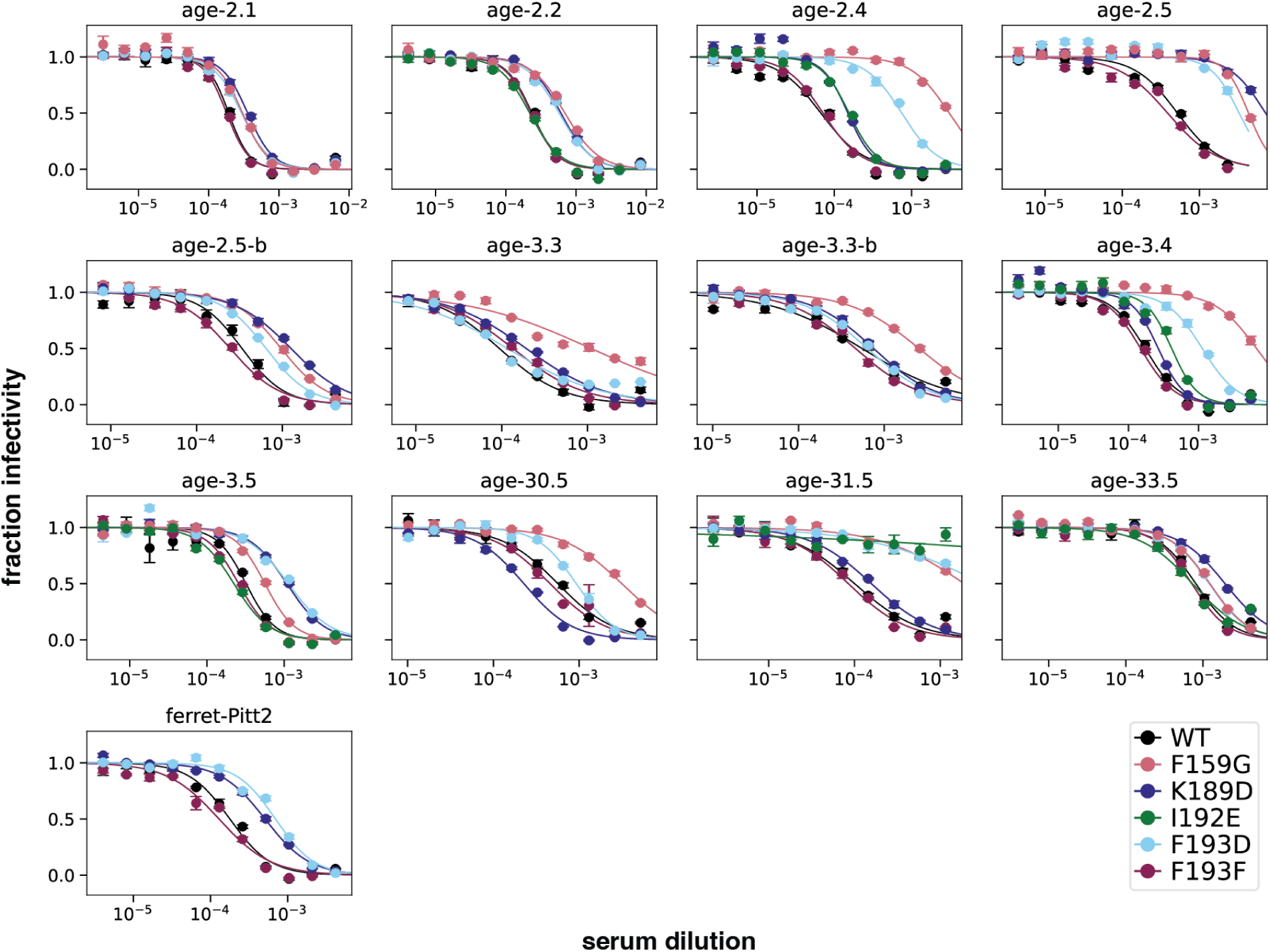
Neutralization assay results for validation of deep mutational scanning measurements against the A/Perth/16/2009 library. Neutralization curves are shown for unmutated A/Perth/16/2009 H3 HA (‘WT’) and selected mutants against all human sera from the 2010-2011 cohort, plus one representative ferret serum.

**Table S1.** Age, serum collection date, recent vaccination history, and relevant medical history for the unvaccinated infant and individuals in the 2-5 and 15-20 year age cohorts from Seattle, Washington.

| Study ID | Age at serum collection (in years) | Serum collection date (quarter) | Flu shot 1 date (quarter) | Flu shot 2 date (quarter) | Flu shot 3 date (quarter) | Immunocompromised | Recent intravenous IgG treatment? | Infection history |
| --- | --- | --- | --- | --- | --- | --- | --- | --- |
| 2462 | 0.2 | Apr-June 2020 | None | None | None | No | No | unknown |
| 2323 | 3 | Apr-June 2020 | Oct-Dec 2019 | Oct-Dec 2018 | Oct-Dec 2017 | No | No | unknown |
| 2367 | 3 | Apr-June 2020 | Jul-Sep 2019 | Oct-Dec 2018 | Jan-Mar 2018 | No | No | unknown |
| 2388 | 4 | Apr-June 2020 | Oct-Dec 2019 | Oct-Dec 2017 | Jan-Mar 2017 | No | No | unknown |
| 2389 | 4 | Apr-June 2020 | Oct-Dec 2019 | Oct-Dec 2018 | Jul-Sep 2017 | No | No | unknown |
| 3944 | 4 | Apr-June 2020 | Oct-Dec 2019 | Oct-Dec 2017 | Jan-Feb 2017 | No | No | unknown |
| 3973 | 2 | Apr-June 2020 | Oct-Dec 2019 | Jan-Mar 2019 | Oct-Dec 2018 | Yes | No | unknown |
| 4584 | 2 | Apr-June 2020 | Oct-Dec 2019 | Jan-Mar 2019 | Jan-Mar 2018 | No | No | unknown |
| 4299 | 4 | Apr-June 2020 | Oct-Dec 2019 | Oct-Dec 2018 | Oct-Dec 2017 | Yes | No | unknown |
| 3862 | 15 | Apr-June 2020 | None | None | None | No | No | unknown |
| 2350 | 20 | Apr-June 2020 | Oct-Dec 2019 | unknown | unknown | Yes | No | unknown |
| 2365 | 17 | Apr-June 2020 | Oct-Dec 2018 | Oct-Dec 2016 | Jul-Sep 2013 | No | No | unknown |
| 2380 | 15 | Apr-June 2020 | Oct-Dec 2019 | Oct-Dec 2018 | Oct-Dec 2017 | No | No | unknown |
| 2382 | 19 | Apr-June 2020 | Jul-Sep 2018 | Oct-Dec 2017 | Jul-Sep 2015 | No | No | unknown |
| 3856 | 20 | Apr-June 2020 | Oct-Dec 2019 | Oct-Dec 2012 | None | Yes | No | Influenza B positive in Jan-Mar 2020 |
| 3857 | >15 years | Apr-June 2020 | unknown | unknown | unknown | unknown | No | unknown |
| 3866 | 18 | Apr-June 2020 | Oct-Dec 2018 | Oct-Dec 2017 | Oct-Dec 2016 | Yes | Yes | unknown |

**Table S2.** Age, serum collection date, and recent vaccination history for individuals in the 40-45 year age cohort from Seattle, Washington.

| Study ID | Age at serum collection (in years) | Serum collection date (quarter) | Received flu vaccine in 2019-2020 season? | 2019-2020 flu vaccine date (quarter) | Received flu vaccine in 2018-2019 season? |
| --- | --- | --- | --- | --- | --- |
| 34C | 45 | Apr-June 2020 | Yes | Oct-Dec 2019 | Yes |
| 199C | 44 | Apr-June 2020 | Yes | Oct-Dec 2019 | Yes |
| 197C | 42 | Jul-Sep 2020 | No | N/A | Yes |
| 18C | 43 | Jul-Sep 2020 | Yes | Jul-Sep 2019 | Yes |
| 33C | 43 | Jul-Sep 2020 | Yes | Oct-Dec 2019 | Yes |
| 215C | 43 | Jul-Sep 2020 | Yes | Oct-Dec 2019 | Yes |
| 74C | 42 | Oct-Dec 2020 | No | N/A | No |
| 210C | 45 | Oct-Dec 2020 | Yes | Oct-Dec 2019 | Yes |
| 150C | 43 | Oct-Dec 2020 | Yes | Oct-Dec 2019 | Yes |
| 68C | 44 | Oct-Dec 2020 | No | N/A | Yes |

**Table S3.**
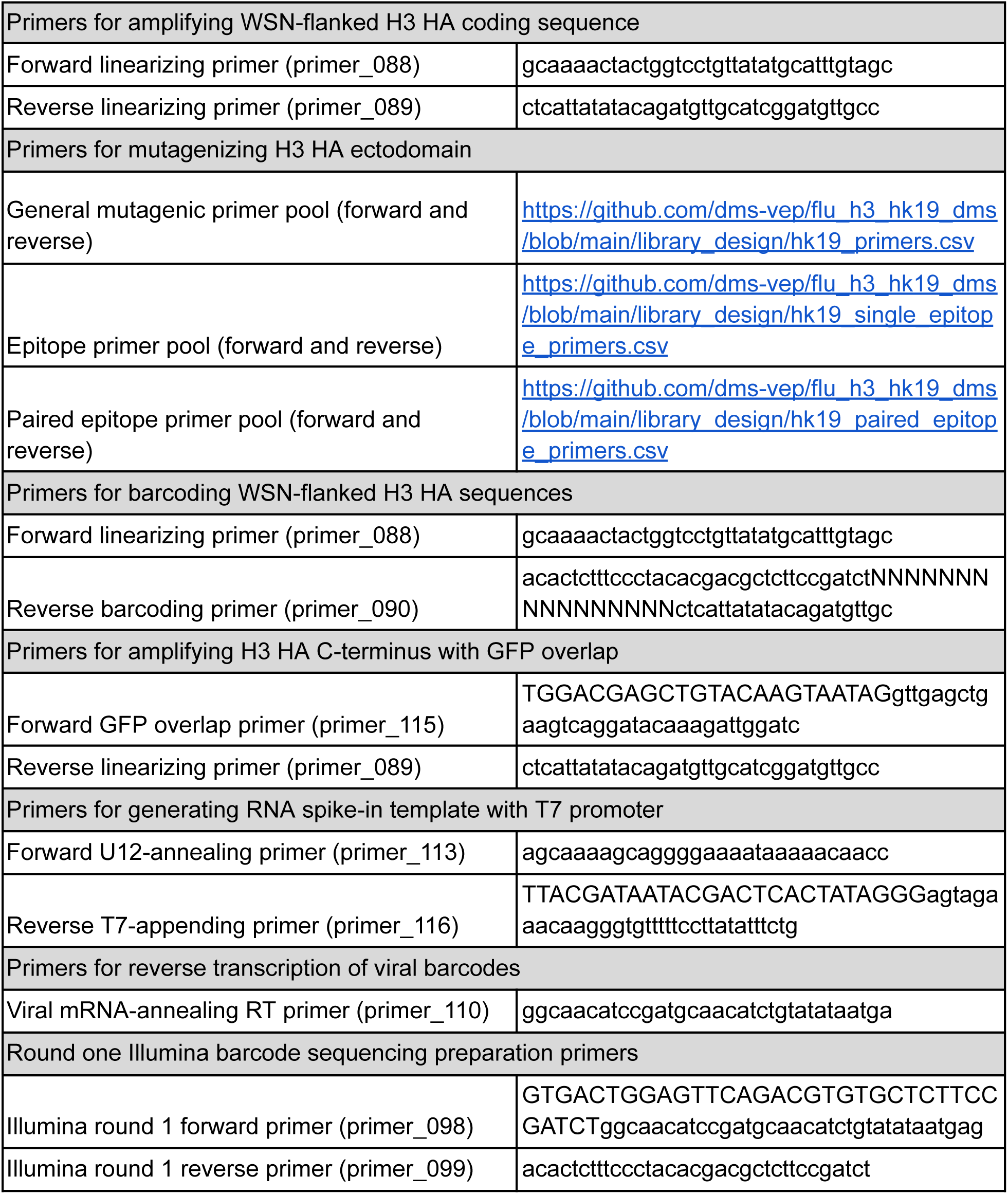
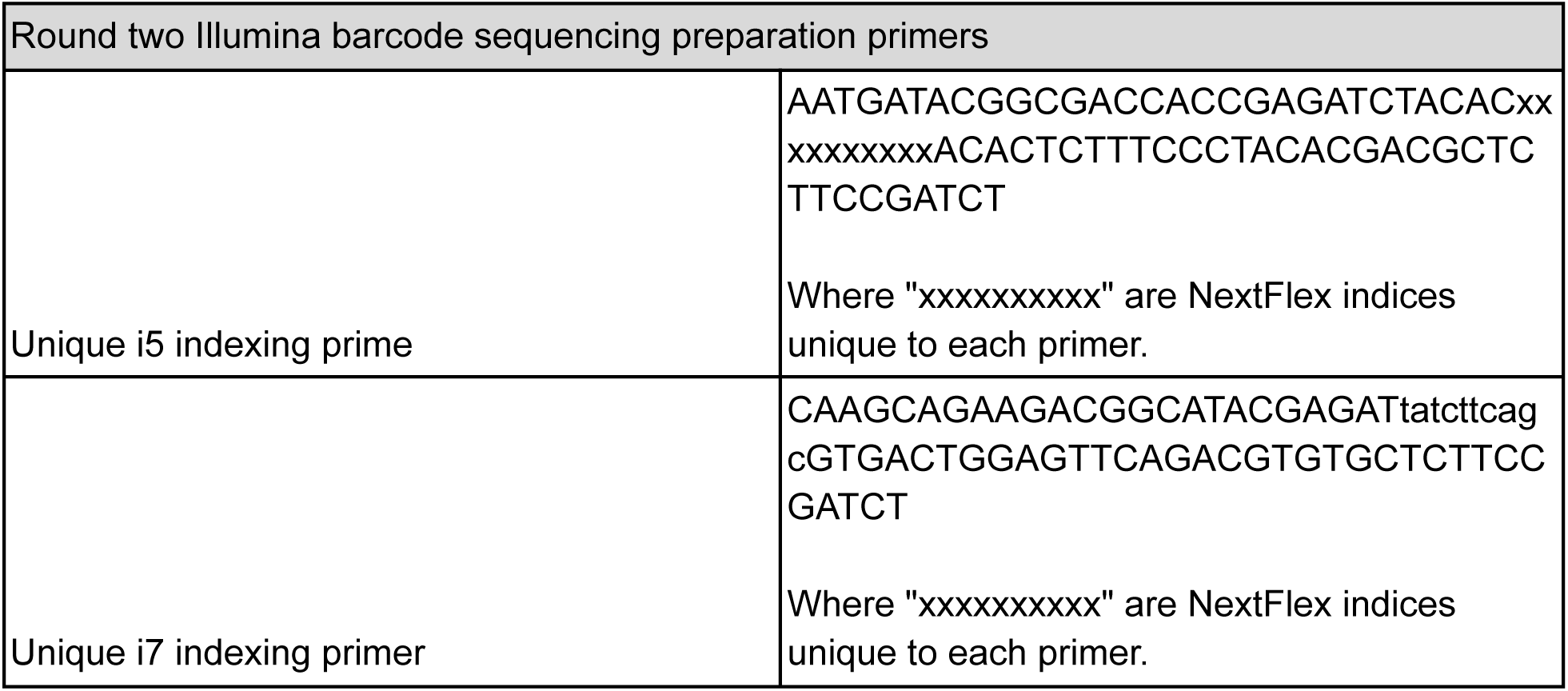
Primers and sequences referenced in methods.

